# Topological Sholl Descriptors For Neuronal Clustering and Classification

**DOI:** 10.1101/2021.01.15.426800

**Authors:** Reem Khalil, Sadok Kallel, Ahmad Farhat, Paweł Dłotko

## Abstract

**Motivation:** Given that neuronal morphology can widely vary among cell classes, brain regions, and animal species, accurate quantitative descriptions allowing classification of large sets of neurons is essential for their structural and functional characterization. However, robust and unbiased computational methods currently used to characterize groups of neurons are scarce.

**Results:** In this work, we introduce a novel and powerful technique to study neuronal morphologies. We develop mathematical descriptors that quantitatively characterize structural differences among neuronal cell types and thus allow for their accurate classification. Each Sholl descriptor that is assigned to a neuron is a function of a distance from the soma with values in real numbers or more general metric spaces. To illustrate the use of Sholl descriptors, six datasets were retrieved from the large public repository http://neuromorpho.org/ comprising neuronal reconstructions from different species and brain regions. Sholl descriptors were subsequently computed, and standard clustering methods enhanced with detection and metric learning algorithms were then used to objectively cluster and classify each dataset. Importantly, our descriptors outperformed conventional techniques and thus provide a practical and effective approach to the classification of diverse neuronal cell types, with the potential for discovery of subclasses of neurons.

## 1. Introduction

Neuronal morphology dictates how information is processed within neurons [1], as well as how neurons communicate within networks [2]. Thus, given the large diversity in dendritic morphology within and across cell classes, quantifying variations in morphology becomes fundamental to elucidate neuronal function. The two major classes of neurons in the neocortex are principal cells (pyramidal cells), and GABAergic interneurons. Pyramidal cells play a critical role in circuit structure and function, and are the most abundant type in the cerebral cortex (70-80% of the total neuronal population) [3]. The morphology of pyramidal cells can vary substantially among cortical areas within a species [4, 5, 6, 7], and across species [8, 9]. Similarly, neocortical GABAergic interneurons are important in shaping cortical circuits, accounting for 10-30% of all cortical neurons [10, 11]. Classification of GABAergic interneurons has proved to be especially challenging due to their diverse morphological, electrophysiological, and molecular properties [12, 13]. Importantly, morphological differences among classes and subclasses of pyramidal cells and interneurons are presumed to be functionally relevant. Moreover, changes in neuronal morphology is thought to underlie various neurodevelopmental [14], and acquired [15, 16, 17, 18, 19] disorders. Thus, given the key role of pyramidal cells and interneurons in cortical function in health and disease, it is important to differentiate among their subclasses through rigorous descriptors and classification tools.

Standard neuronal descriptors rely on measurements of morphological features typically acquired from digital neuronal reconstructions. Feature measurements are subsequently used to quantitatively assess and cluster cell classes [20], using standard supervised [13], and unsupervised [21, 22, 23] clustering algorithms. The raw morphological desciptors, provided by standard methods often fail to discriminate among neuronal classes that are visually very different (§3). Therefore, there is a demand for more robust and general methods for discriminating among diverse neuronal cell types and larger datasets. In recent years, the field of computational topology has become increasingly more popular in the characterization of tree structures, including neurons. For example, [24] developed a new algorithm called ‘Topological morphological descriptor’, TMD which is inspired by topological data analysis (i.e. persistence diagrams) to classify families of neurons. In a more recent study by [25], the authors use TMD to classify cortical pyramidal cells in rat somatosensory cortex. The topological classification was largely in agreement with previously published expert assigned cell types. Furthermore, [26] present a framework based on persistence homology to compare and classify groups of neurons. Despite their partial success, the available methods fail to fully capture the subtle morphological differences among families of neurons as we illustrate in the present work.

Here we take a novel approach to the classification of neurons. We view a morphological feature describing a neuron as a function which takes values in the real numbers, or more generally in some relevant metric space, and varies as a function of distance from the soma. We choose neurobiologically meaningful features which are known to differ among cell classes, such as branching pattern, tortuosity, and taper rate. Each feature gives rise to what we refer to as a *Sholl descriptor*. This Sholl descriptor is a rule that assigns a metric element, such as a number or a persistence diagram, to every given neuron and at any given distance from its soma. This assignment is an isometry invariant which means that it does not depend on how the neuron is placed in space, as long as its soma is at the origin. Moreover, all value assignments are *stable* which implies that they are close in values whenever the neurons are sufficiently close in a piecewise differentiable way (Supplementary §2). The construction we just outlined endows the set of neurons up to isometry with a *descriptor metric* for every Sholl descriptor. Our approach is useful in that every morphological feature turns the set of isometric neurons into a metric space. The closer the neurons are in the underlying descriptor metric, the more of this feature they share. This method gives a powerful, objective, and interpretable tool to compare and analyze neuronal morphologies.

In the course of our work, we developed eight Sholl descriptors, representing eight features which we then use in both unsupervised and supervised settings to cluster and classify families of neurons. Using diverse datasets, we identify key descriptors that reveal differences and similarities between neuronal classes. Our discrimination results were significantly better in separating different neuronal cell types than clustering methods based on raw quantification of features. Certain descriptors result in complete separation of selected groups of neurons. Thus, our highly effective and powerful classification tool could be used for the identification of new neuronal cell types, ultimately enhancing our understanding of the morphological diversity and function of neurons in the brain.

## 2. Methods

In this study, we developed a toolkit of descriptors, and proposed their implementation. We introduced an unsupervised clustering and supervised classification framework based on relevant morphological features to differentiate among and classify different neuronal cell types. Our pipeline works as follows: Sholl descriptors are computed on a dataset of neuronal reconstructions. They are then used as an input to a wide range of methods; starting from dendrogram analysis, computations of detection rates (Supplementary §1.4), or more advanced machine learning techniques (Fig. 1). Given a topological or morphological feature of neurons, denoted by the Greek letter *ϕ*, such as branching pattern, tortuosity, total wiring, etc, we associate to such a feature a “Sholl descriptor”. Specifically, a Sholl descriptor is a map from the set of neurons to a metric space of functions that is invariant under certain symmetries and has stability properties (Definition 2.3). We then construct a metric *d*_*ϕ*_ on the set of descriptors of all neurons §2 and use it as a proxy of a distance between those neurons. In other words, we endow the set of neurons, up to isometry, with a metric space structure for every feature *ϕ*. This (Sholl) metric measures proximity of neurons with respect to the particular feature captured by *ϕ*. In particular, the closer the neurons are under that metric, the more of the features captured by *ϕ* they share. The fundamental idea of a Sholl descriptor is presented in Fig. 2. In this case, a branching pattern function is presented for a simple tree. The value of the descriptor for that neuron is the step function on the right.

**Fig. 1.**
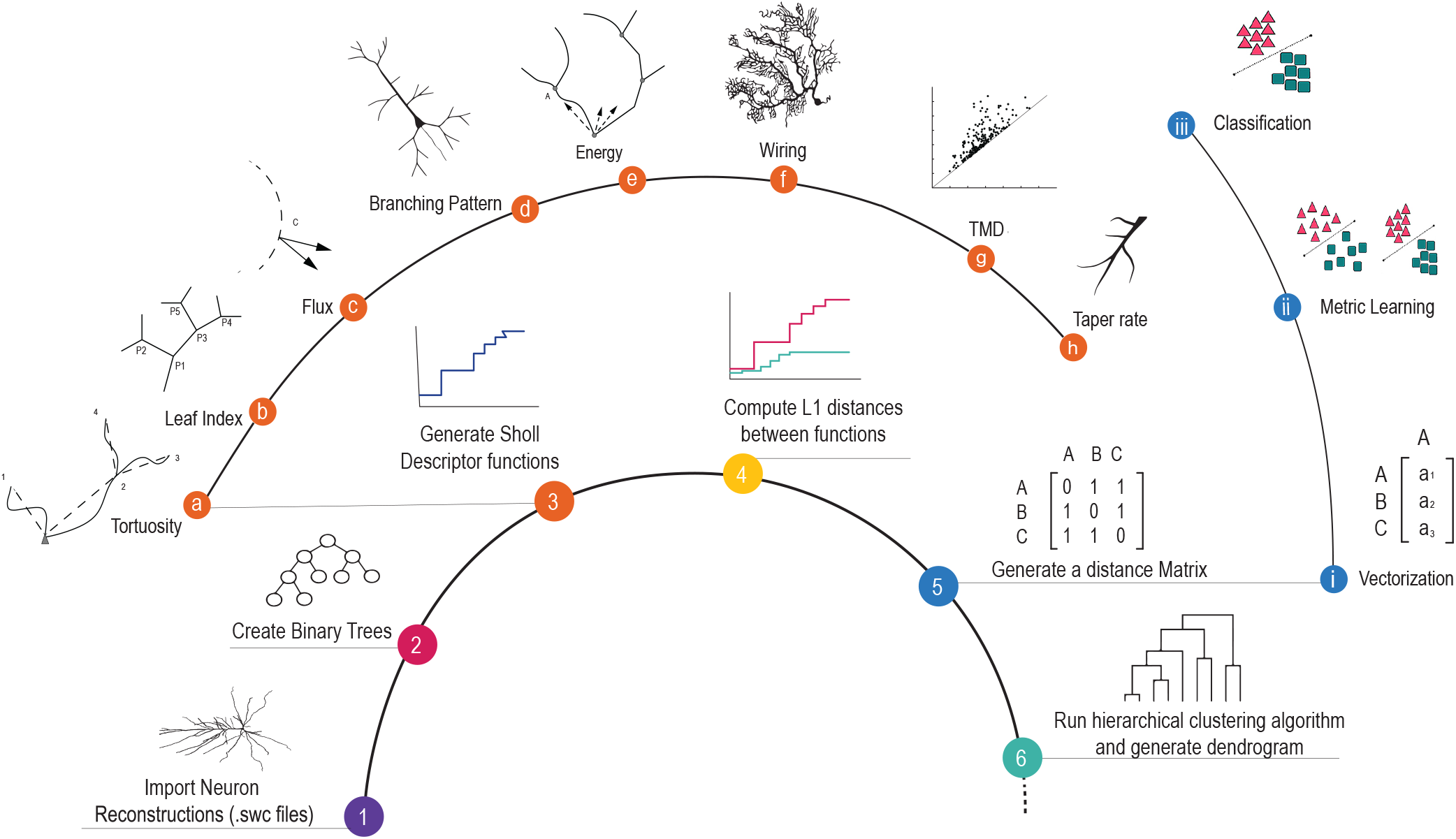
A pipeline to generate and cluster morphological data. Reconstructions of neuronal cell types are encoded in Sholl descriptors which are then used for clustering and classification. A representation of each Sholl descriptor is depicted in steps **a-h**. The process of clustering different neuronal cell types is shown in steps 1-6. Once neurons are vectorized based on descriptor metrics, we apply metric learning techniques to obtain classification (steps **i-iii)**.

**Fig. 2.**
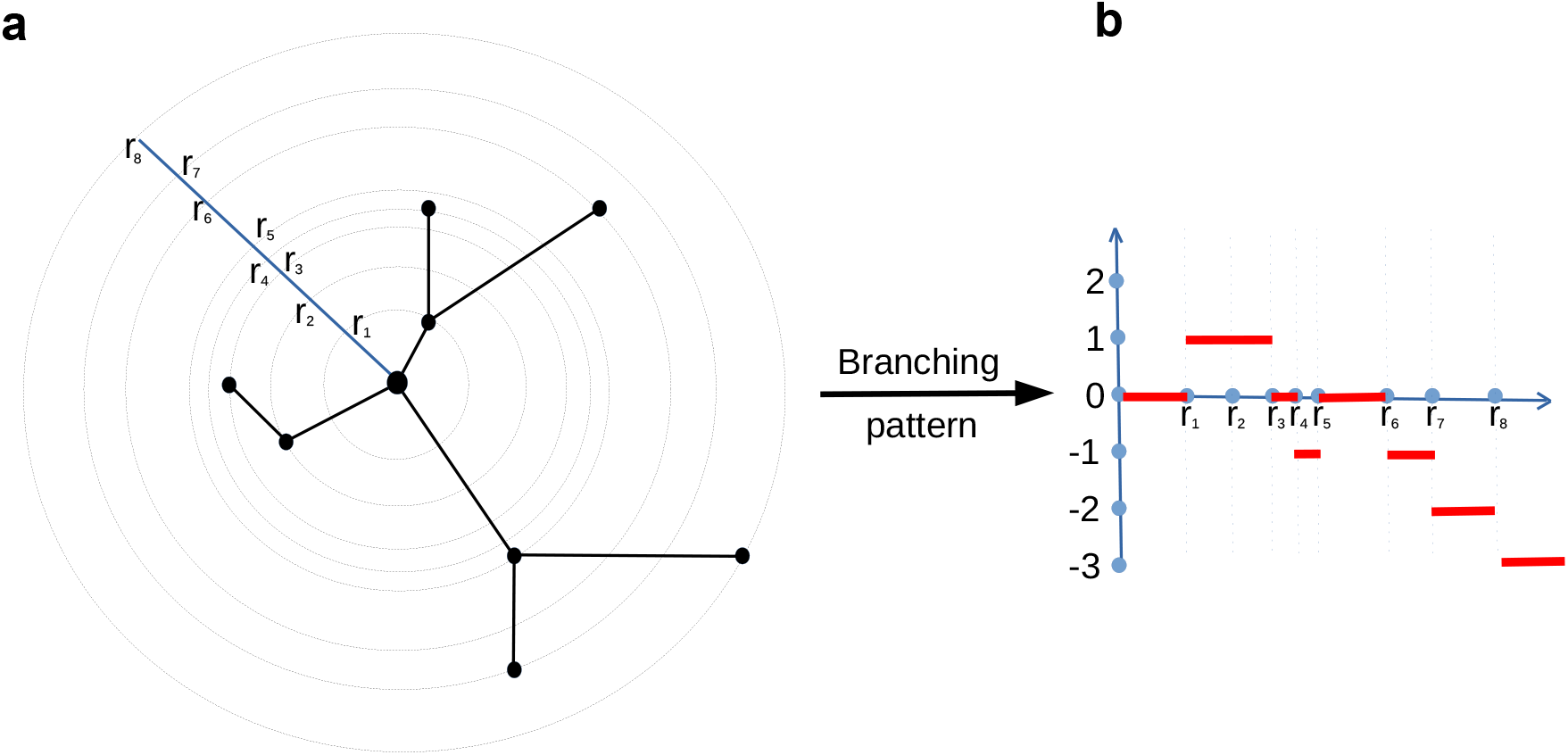
Sholl descriptor using branching pattern. (see Supplementary §1.6.1). (**a**) A representation of a tree. (**b**) the corresponding branching function as a function of distance from soma.

Different descriptors can then be combined to capture multiple aspects of neuron morphology to optimize clustering and enhance classification. We present three different methods of combining the descriptors, each with its own merits. The first combination method is unsupervised in that it does not depend on the way we subdivide our dataset into classes (Supplementary §4.1). It is based on a certain vectorization of our Sholl descriptor that can subsequently be used in unsupervised Machine Learning algorithms. The second method combines distances between multiple features into a new metric by optimizing coefficients of a linear combination of distances using a grid-search. The aim is to maximize separation of classes (Supplementary §4.2). If no such combination can be produced, the classes are indistinguishable under all used morphological descriptors. The third combination method is a supervised classification technique which is achieved by means of metric learning. It works by first pre-selecting features that are relevant for classification based on their detection rates (Supplementary §1.4). This optional step allowed us to enhanced the results of the subsequent classification. Subsequently selected relevant descriptors of neurons are vectorized and a metric learning algorithm is applied to those vectors. It produces an optimal metric that can be subsequently used for classification. The second and third methods are checked for ‘overfitting’ (Supplementary §4.4). The new metric effectively reveals how close in features a random neuron is to the given classes.

In addition to conventional methods, we introduce the notion of “detection” whereby a class *C* is said to be detected 100% by a descriptor *ϕ* if the entire class fits in a ball under the descriptor metric, and no other neuron from any other class is within that ball. Detection is a convenient tool that helps reveal, via a rapid and direct use of our constructed metrics, how much a given feature is able to single out the class *C* from all other considered classes. Detection rates (Supplementary §1.4) are correlated with the corresponding dendrogram obtained via cluster analysis, so a high detection rate leads to neurons in the class clustering together in the accompanying dendrogram.

### 2.1. Representation of neurons

We model a neuron *N* as a tree embedded in 3-dimensional space ℝ^3^. The tree is composed of a collection of rooted binary trees, all having a common root (the soma)^1^. We assume that the root is located at the origin in the considered coordinate system. These rooted treed are also called the “primary” trees.

All 3D neuronal reconstructions used in this paper are acquired from the public repository NeuroMorpho.org [27]. The morphological structure of individual neurons is retrieved from an SWC file which contains a digital representation of the neuron as a tree structure that consists of points in ℝ^3^ joined by edges. Each marker has associated properties such as 1) its 3D spatial coordinates, 2) its radius denoting the thickness of the branch segment at a specific 3D location 3) a node type indicating whether it is soma, axon or dendrite, and 4) one parent marker to which it directly connects through neuronal arbors. The soma is always located at the origin of a reference frame.

It should be noted that well-defined geometric descriptors on neurons, that we are providing in this paper, need to be invariant under the affine isometry group of ℝ^3^. Under the assumptions about representation of a neuron listed above, it is sufficient to only consider invariance by rotations and reflections, disregard translational invariance.

### 2.2. Notation and Terminology

- Capital letter *N* represents a neuron seen as a tree in 3-space.
- A class *C* of neurons is a set containing a selection of neurons of a particular type.
- A node in a neuron *N* is either the soma, bifurcation point, or a termination point(Fig. 3)

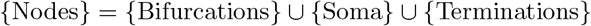 A branchpoint can be used interchangeably with bifurcation point. A leaf can be used interchangeably with a termination point. Note that points of degree two are not explicitly used in the following constructions and hence not considered as nodes.
- The number of terminal nodes in a tree is denoted as degree which is a proxy for tree complexity. The number of branches of a tree is twice the degree of that tree minus one, while the number of bifurcations is degree minus 1.
- Radial distance means Euclidean distance as measured from a point to the soma, see Fig. 3.
- Path distance is the distance from a point to the soma along a dendrite, see Fig. 3.
- A branch is a part of the dendrite that lies between two branchpoints or between one branchpoint and a termination point.
- Two nodes are parent-child related if they are adjacent on a branch. The node closer to the soma in path distance is called the parent, and the node farther away from the soma is called the child.
- A neuron has span *R*(*N*) if it can fit in a ball centered at the soma of radius *R*(*N*), and in no smaller ball. In practice and almost always, *R*(*N*) is the largest radial distance from soma to any of the nodes.
- *L*(*N*) is the length of the longest dendrite stemming from soma and ending at a termination point.
- A neuronal feature is denoted by the Greek letter *ϕ*. It is always topological or morphological in nature, and its associated (Sholl) descriptor will also be denoted by *ϕ*.
- A feature *ϕ* gives rise to a metric on the set of neurons which is denoted by *d*_*ϕ*_.

**Fig. 3.**
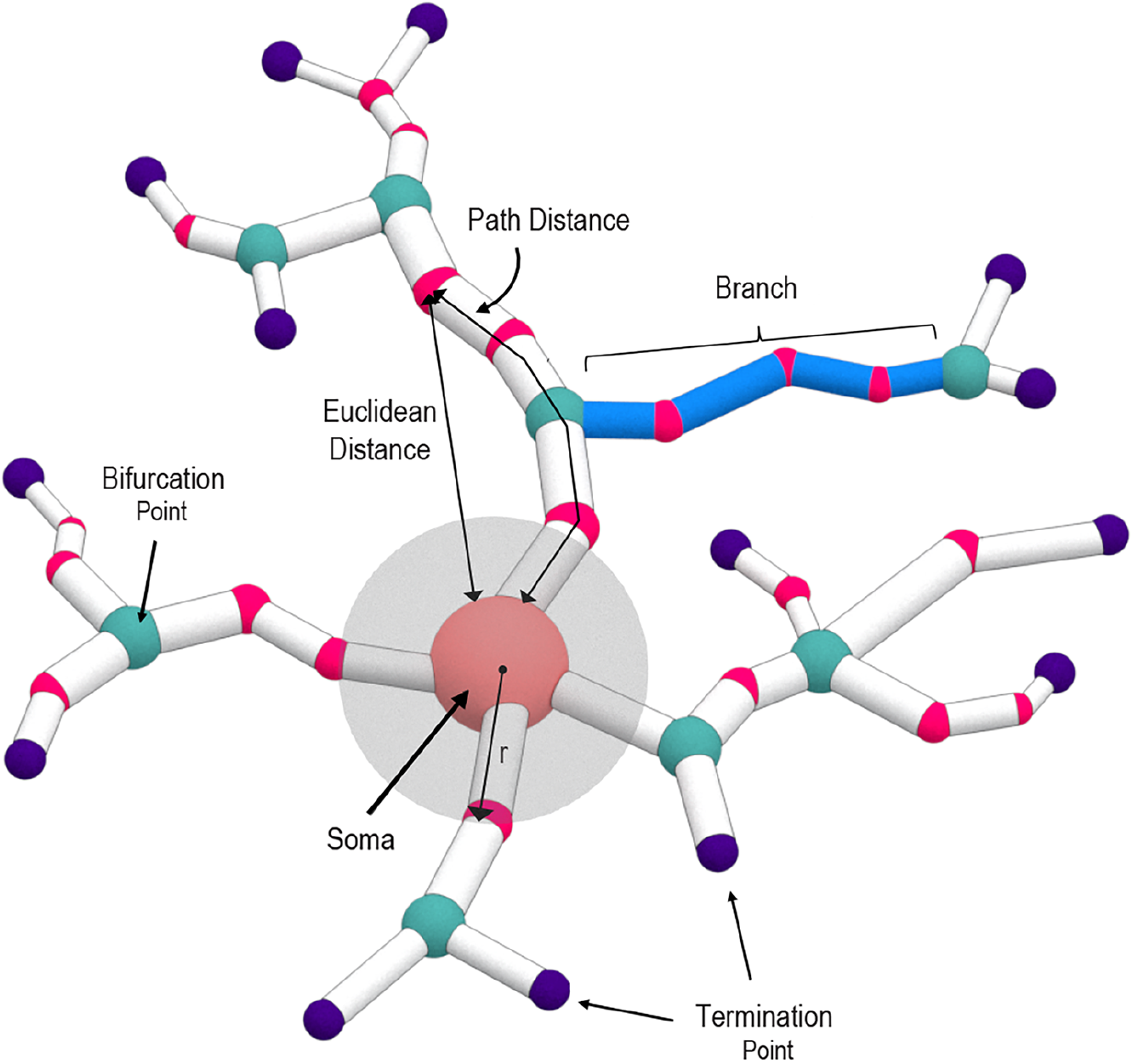
Representation of a neuronal tree structure. Schematic of a neuron illustrating key definitions

### 2.3. Sholl descriptors

Below we detail the construction and definition of Sholl descriptors.

#### Definition 2.1.

*A “Sholl descriptor” is any rule that associates to a given neuron (represented as a tree embedded in* ℝ^3^*) a compactly supported function whose independent variable is either path or radial distance from the soma, and whose values are in a metric space X. We further require this function to be both isometry invariant and stable with respect to reconstruction errors*.

More precisely a Sholl descriptor 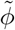 associates a function 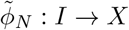 to a given neuron *N* ⊂ ℝ^3^. Here *I* is either the interval [0, *R*(*N*)] or [0, *L*(*N*)] (see §2.2 for definitions) and *X* is a metric space. The *ϕ* is stable in the sense of Supplementary 2. Isometry invariance means that if *N′* is obtained from *N* by rotation or reflection by a plane passing through the soma, then 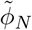 and 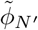 are identical functions.

Since we want our constructions to be independent of scale, the domains of all functions are normalized to an interval [0, 1] (see Supplementary §1.1). Effectively, *a Sholl descriptor associates to every neuron N a Sholl function* [0, 1] → *X, with X a metric space, satisfying the stability and isometry invariance properties*.

The following are Sholl descriptors that we discuss in this paper (mathematical details are given in §2.3).

1. **Branching pattern**: This is an integer–valued descriptor equals the number of bifurcations minus the number of leaves in a distnace *r* from the soma. As the radius *r* changes, this number evolves. A different Sholl descriptor is obtained when a radial distance is replaced with a path distance from the soma. In this case, the connected component of a tree inside a radius *r* ball centered in the soma containing the soma is considered. The branching pattern descriptor quantifies dendritic complexity or branching structure, and the distribution of dendritic branches. The branching structure of a neuron is important for determining function. For example, in primary visual cortex, branching structure is related to orientation and direction selectivity of neurons.
2. **Tortuosity**: Tortuosity of a given path is defined as the ratio of the path length by the Euclidean distance of its endpoints. The tortuosity descriptor at a distance *r* from the soma measures the mean tortuosity of all branches connected to the soma within the sphere of a radius *r* centered at the soma. This descriptor informs about the dendrite’s growth mechanisms and its method of reaching synaptic targets.
3. **Flux**: This descriptor starts with a sphere of a radius *r* centered at the soma. For all the dendrites intersecting that sphere it associates to *r* the sum of the angles between dendrites and normal directions to the sphere. The flux construction is related to, and can be viewed as an extension of the root-angle construction in [44]. This construction associates to a leaf the angle between the radial normal and the main branch ending at that leaf. This measure quantifies centripetal bias which is based on the branching characteristics of dendrites.
4. **Taper Rate**: This construction is based on the width of dendritic segments at bifurcations, taken as a function of path distance to the soma. To be precise, the branching nodes are ordered according to their path distance to the soma. The value of the Taper Rate, for a radius *r*, is equal to the dendritic diameter at the closest node in a distance at most *r* from the soma (see Supplementary §1.6.3). Dendritic tapering is a measure of the change in width along a dendritic segment from node to node. Dendritic tapering towards the terminal tips maximizes current transfer and thus leads to enhanced integration of synaptic inputs at different locations.
5. **Leaf Index**: This construction counts the number of leafs that can be reached from a given node. If the node is taken to be the soma, this number is the total number of leaves in the tree. If the node is taken to be a leaf, the value is one (that leaf). The Leaf Index considers branching nodes as a function of radial distance. For a given value *r*, consider the last branchpoint *b* that is not farther away from soma than *r*. The value of Leaf Index at the radius *r* is equal to the number of leafs reachable from *b*. This measure quantifies centripetal bias which is based on the branching characteristics of dendrites
6. **Energy**. This descriptor is based on the idea of viewing the nodes as charged electrons that generate electric fields. Using the superposition principle, the resulting vector field at the soma is recorded. Its length is an isometry invariant quantity. For any given radius *r*, we can carry out the same construction using only those nodes not further away from soma than *r*, and take the sum of their contributions at the soma (again only the length is measured). This descriptor gives a measure of how the nodes are distributed relative to the soma. If we divide space in octants, with the soma at the origin, then the more nodes present in the same octant, the greater the energy. This is similar to caulescence which refers to a prominence of a main path in a tree structure to reveal tree asymmetry [28]. High caulescence values reflects dendrites that follow a similar route while low values imply a more balanced tree structure without a distinct main path.
7. **Total Wiring**. Take a tree being a connected component of a neuron that contain soma, restricted to a ball of a radius *r*. This construction measures the total path length of this tree. This is the sum of the path distances of all dendritic segments. Greater total wiring dendritic length means greater invaded space which render the dendrites available to more incoming connections and greater connectivity.
8. **TMD**. This is the Topological Morphological Descriptor of [24] redesigned to be a Sholl function. The metric space target in this case is the space of persistent diagrams with the Wasserstein metric.

## 3. Results

We propose eight descriptors based on the morphological features listed above. A schematic of each descriptor is illustrated in steps a-h in Fig. 1, and all terms are defined in §2.3. We illustrated the discriminative accuracy of our Sholl descriptors on six datasets, providing evidence that relevant Sholl descriptors can reliably discriminate among different classes of neurons in agreement with previously published assignment. All datasets were downloaded from Neuromorpho.org [27]. They were chosen to cover diverse types and subtypes of neurons across different regions and animal species. For convenience, in the discussion below, we say a class has been *detected* by a descriptor *ϕ* if the rate of detection for that class is at least 90% (see discussion prior to §2.1, and details in §1.4).

### L-measure versus Sholl descriptors

As a proof of concept, we applied our Sholl descriptors on **Dataset 1** which comprised three different neuronal cell types in the mouse brain: retinal ganglion cells (n=10), cerebellar purkinje cells (n=9), and interneurons in the medial prefrontal cortex (n=10). The aim was to choose strikingly different neuronal cell types that could easily be clustered. A representative neuron from each type is shown in Fig. 5a-c. Seven different Sholl functions were computed on this set (all but taper-rate since the information about diameter of dendrites was not available) and distance matrices between neurons have been computed for each Sholl function. As shown in the detection Table 1, the performance of all descriptors was optimal in that each class was completely detected by at least two descriptors. Remarkably, the branching pattern descriptor detected all three classes, suggesting that this descriptor alone is sufficient in classifying this particular dataset. The parameterized TMD descriptor (TMD Sholl) performed equally well as other descriptors, and better than its classical version. The dendrogram based on the TMD Sholl is shown in Fig. 5f. The horizontal axis of the dendrogram represents all the neurons in this dataset while the vertical axis represents the distance between clusters. Interestingly, interneurons (Fig. 5b) were fully detected by every single descriptor, suggesting that this cell type is characterized by a unique set of morphological features.

**Table 1.** Detection rates applied to all datasets based on all descriptors. Numbers represent percentages. Green represents complete detection of a class (100%), while pink represents detection rates between 80%-100%. The taper rate was only run on Dataset 2 since measurements of dendritic width were available for this dataset.

| Dataset | Description | Energy | Flux | Leaf index | Branching | Wiring | Tortuosity | TMD Classical | TMD Sholl | Taper rate |
| --- | --- | --- | --- | --- | --- | --- | --- | --- | --- | --- |
| 1 | Mouse retinal ganglion | 91 | 90 | 100 | 100 | 82 | 90 | 91 | 91 |  |
| . | Mouse cerebellum Purkinje | 89 | 82 | 31 | 100 | 78 | 73 | 90 | 100 |  |
| . | Mouse L5 interneuron, mpfc | 100 | 100 | 100 | 100 | 100 | 100 | 100 | 100 |  |
| 2 | Mouse L5 Martinotti, occipital | 48 | 73 | 68 | 68 | 73 | 91 | 64 | 55 | 43 |
| . | Mouse medium spiny, basal ganglia | 40 | 67 | 59 | 60 | 60 | 63 | 47 | 53 | 87 |
| . | Mouse retinal ganglion | 100 | 90 | 82 | 90 | 90 | 64 | 91 | 70 | 60 |
| . | Mouse pyramidal, hippocampus | 62 | 73 | 50 | 64 | 50 | 92 | 50 | 83 | 82 |
| . | Mouse stellate, somatosensory | 60 | 36 | 43 | 33 | 25 | 30 | 33 | 33 | 67 |
| 3 | Monkey L3 pyramidal, V1 | 60 | 91 | 70 | 100 | 91 | 70 | 90 | 91 |  |
| . | Monkey L3 pyramidal, V2 | 64 | 82 | 53 | 82 | 82 | 60 | 58 | 70 |  |
| . | Monkey L3 pyramidal, V4 | 46 | 82 | 43 | 90 | 91 | 58 | 70 | 82 |  |
| 4 | Rat L5 pyramidal, primary somatosensory | 55 | 100 | 75 | 100 | 90 | 75 | 85 | 95 |  |
| . | Rat L5 pyramidal, secondary motor | 71 | 94 | 48 | 76 | 94 | 71 | 82 | 94 |  |
| . | Rat L5 pyramidal, medial prefrontal | 75 | 100 | 100 | 100 | 84 | 84 | 95 | 95 |  |
| 5 | Rat L5 pyramidal somatosensory (control) | 65 | 67 | 67 | 67 | 65 | 73 | 70 | 67 |  |
| . | Rat L5 pyramidal somatosensory (experimental) | 56 | 56 | 55 | 57 | 50 | 67 | 69 | 63 |  |
| 6 | Mouse cerebellum purkinje | 93 | 80 | 57 | 90 | 57 | 90 | 82 | 91 |  |
| . | Mouse area17 spiny | 41 | 99 | 81 | 95 | 81 | 81 | 94 | 55 |  |
| . | Mouse hippocampal granule | 44 | 80 | 83 | 95 | 84 | 80 | 70 | 68 |  |
| . | Mouse hippocampal microglia | 44 | 62 | 80 | 70 | 66 | 80 | 70 | 72 |  |
| . | Mouse spinal cord interneuron | 35 | 58 | 50 | 55 | 43 | 64 | 51 | 49 |  |

**Fig. 4.**
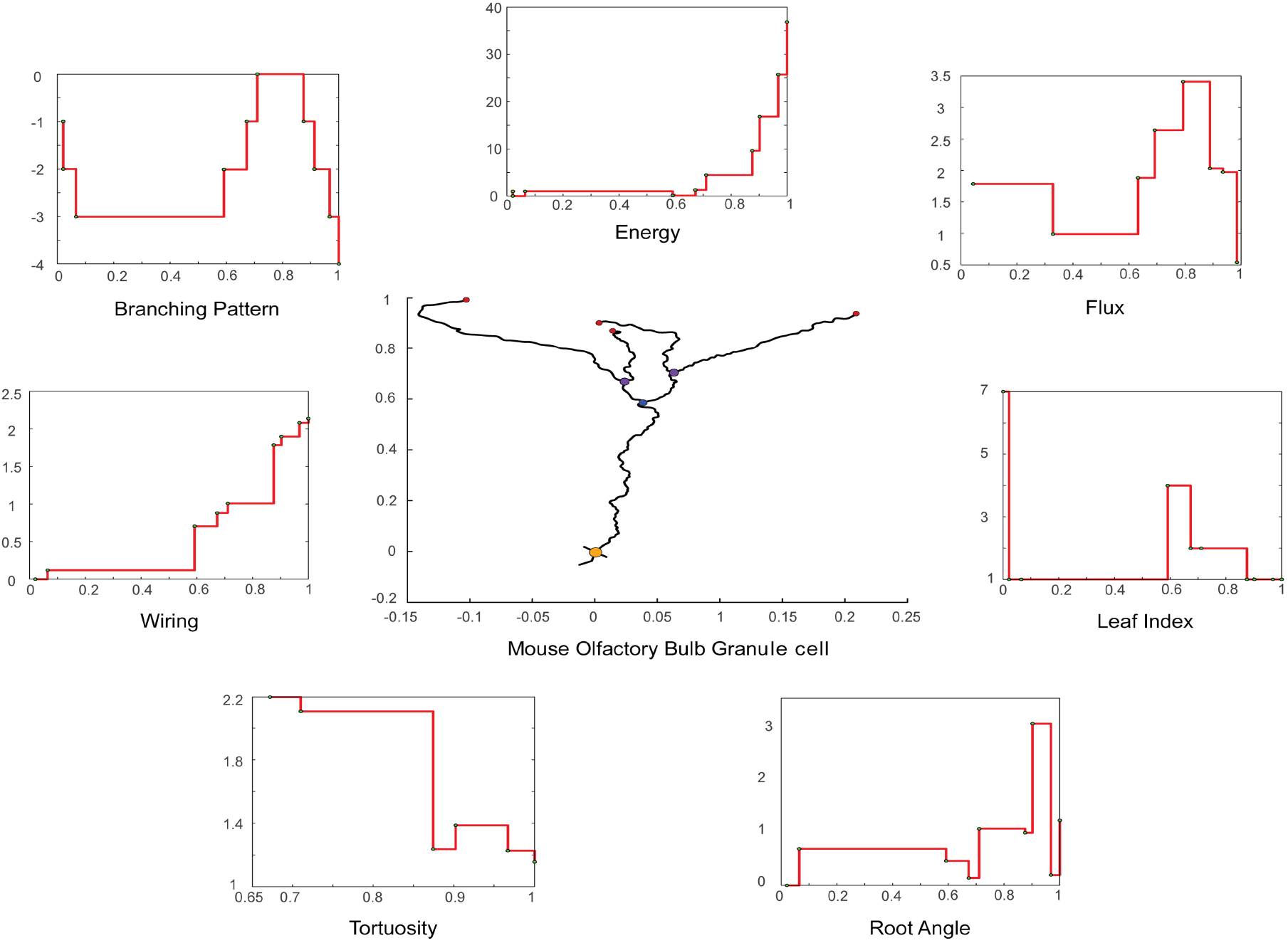
Representative neuronal reconstruction of a granule cell in mouse ol-factory bulb with the corresponding step functions for seven real-valued Sholl descriptors. This neuron was chosen because of its simple branching pattern which consists of 3 bifurcation points and 7 leaves. The step function for the “taper rate” descriptor is missing for lack of data. Also, the “TMD” descriptor is missing here, as it is not real-valued and cannot be easily presented.

**Fig. 5.**
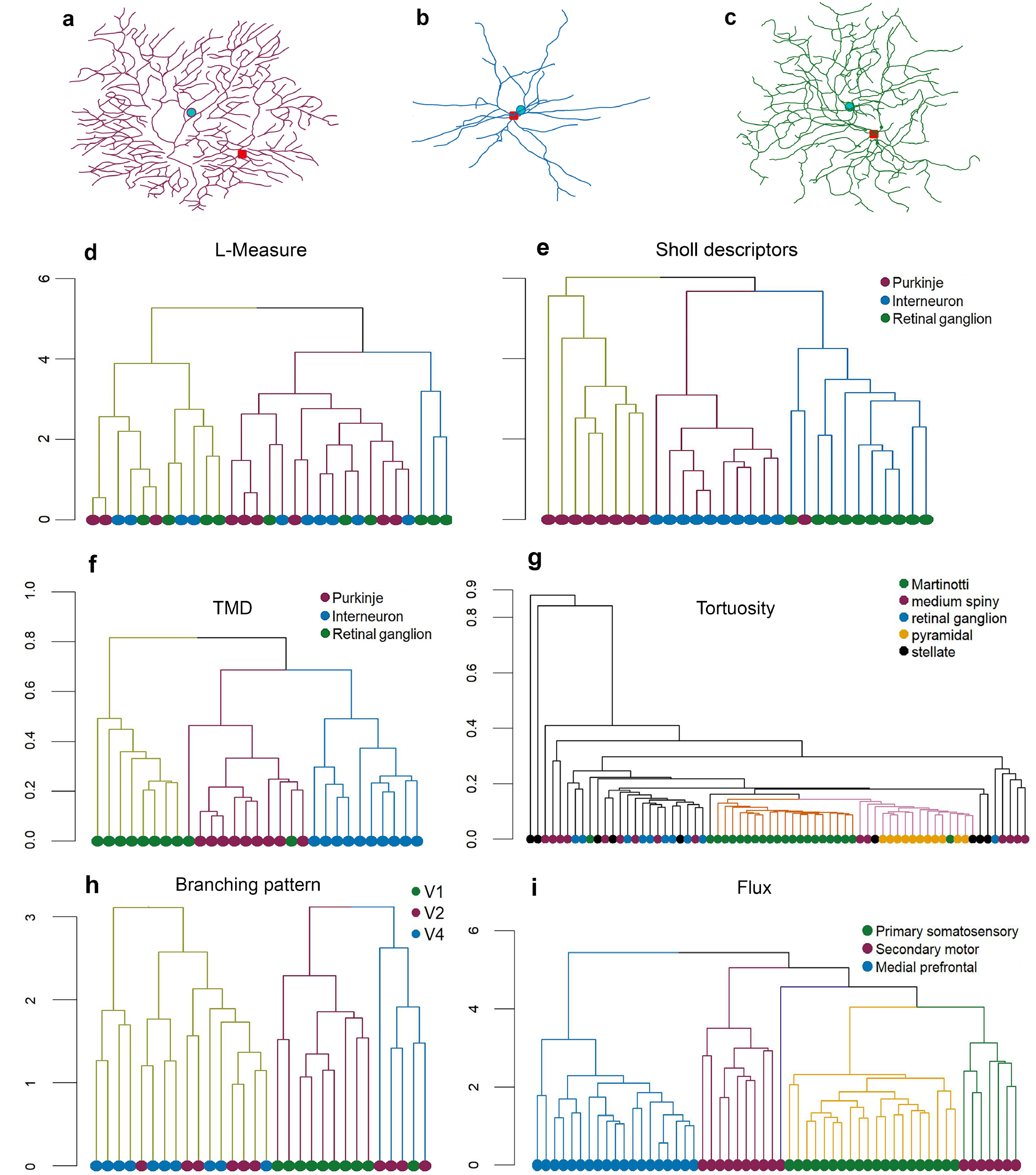
Hierarchical clustering trees for all datasets. Representative neuronal reconstructions of (**a**) purkinje, (**b**) interneuron, and (**c**) retinal ganglion cell, in magenta, blue, and green colors respectively. Representative dendrograms for morphological parameters extracted from (**d**) L-measure software, (**e**) Sholl descriptors. Dendrograms based on descriptors for (**f**) TMD for Dataset 1, (**g**) Tortuosity for Dataset 2, (**h**) Branching pattern for Dataset 3, and (**i**) Flux for Dataset 4. For the reconstructions (**a**,**b**,**c**), the red dot represents the soma and the green dot is the barycenter of all nodes.

The described Sholl descriptors as well as standard L-measures were subsequently used as an input to conventional clustering algorithms. The aim was to determine whether these unsupervised methods will automatically detect the prescribed classes of neurons in the considered dataset and if our Sholl descriptors will do it better than the standard L-measures.

First, for each considered neuron, we combined the morphological parameters from our eight descriptors to a vector representing neuron (as detailed in Supplementary §4.1). A hierarchical cluster analysis algorithm were applied to those vectors. Next, for each considered neuron, we extracted 10 morphological parameters using the L-Measure software [29], combined them into a vector, and applied the same cluster analysis algorithm on it. To make a fair comparison, we chose specific morphological parameters from L-Measure to match features that are captured by our descriptors (number of branches, leaves, bifurcations, contraction, branch angle, taper rate, path distance, and Euclidean distance). Fig. 5 shows a dendrogram of the linkage distances between the 29 neurons based on L-measure extracted features (Fig. 5d) and Sholl descriptors (Fig. 5e). Neurons are color coded according to morphological type. Cluster analysis based on features captured by the Sholl descriptors results in clear separation of classes into three clusters (Fig. 5e), whereas the L-Measure method returns two clusters with significant intermingling of neurons from the three types (Fig. 5d). Therefore, the results demonstrate that our combined descriptors can outperform conventional methods in clustering different neuronal cell types.

### Distinguish among classes based on a single feature

**Dataset 2** included 67 neurons from five different regions of the mouse brain: retinal ganglion cell (n=10), basal ganglia medium spiny (n=15), somatosensory stellate (n=9), hippocampal pyramidal (n=11), somatosensory Martinotti (n=22). This is the only dataset that included dendritic width in the reconstructions, which allowed us to use the taper rate descriptor. The dendrogram in Fig. 5g shows the cluster analysis based on the tortuosity descriptor function. This Sholl descriptor resulted in efficient separation of neuronal cells types, particularly for the Martinotti (detection rate=91%), and hippocampal pyramidal cells (detection rate=92%). Table 1 reports the detection rates (see Supplementary §1.4) for all descriptor functions for Dataset 2. Detection rates highlighted in pink are above 80% while rates of 100% are highlighted in green. In contrary, the leaf index descriptor performed poorly in separating four of the five neuronal cell types, as evidenced by the low detection rates. This suggests that this particular morphological feature is largely uniform across these cell types (Martinotti, medium spiny, pyramidal, and stellate) while others allow for a good separation of the considered classes of neurons.

**Dataset 3** comprises pyramidal cells in layer 3 of different cortical areas of the vervet monkey brain: primary visual cortex (V1) (n=10), V2 (n=10), and V4 (n=10). These reconstructions consisted of only basal dendrites. Prior reports have revealed regional differences in pyramidal cell morphology in the monkey brain [4, 8]. Specifically, pyramidal cell size, dendritic complexity, and spine density increase from primary visual cortex (V1) to higher order visual areas. Therefore, as a proof of concept we sought to recapitulate these findings by running our descriptors on reconstructions of pyramidal neurons from different areas in the visual cortical hierarchy. We expected at least to cluster pyramidal neurons from V1, V2, and V4, based on the branching pattern descriptor. Indeed, the dendrogram in Fig. 5h based on this descriptor reveals excellent separation of V1 neurons with some intermingling among V2 and V4 neurons. The wiring descriptor performed equally well in clustering, with excellent separation of V1 neurons and V4 neurons, and reasonable separation of V2 neurons (Table 1).

### Subclustering within a neuronal class

To ensure sufficient coverage of neurons from different species, we also tested our Sholl descriptors in clustering pyramidal cells from different cortical areas in the rat brain. Therefore, **Dataset 4** consisted of rat pyramidal cells in layer 5 of somatosensory (n=20), secondary motor cortex (n=15), and medial prefrontal cortex (n=19). The discriminative accuracy in separating the three neuron groups with many of the descriptor functions was very high. For example, the cluster analysis based on the flux descriptor shown in the dendrogram in Fig. 5i resulted in nearly perfect clustering. The detection rate was highest in both medial prefrontal and somatosensory cortex (100% detection), followed by secondary motor cortex (94% detection). Likewise, the branching pattern, wiring, and TMD Sholl descriptors performed equally well as shown in Table 1. Remarkably, the combined descriptor approach yielded complete separation with three distinct clusters (Supplementary Fig. 7a) (see Supplementary §4.2). Interestingly, the majority of pyramidal cells in secondary motor cortex formed their own distinct cluster, while several of these cells were clustered with other pyramidal cells in primary somatosensory cortex. This suggests the existence of two subpopulations of pyramidal cells in secondary motor cortex. Indeed, when we visually examined these neurons, we found striking similarities in morphology with pyramidal cells in primary somatosensory cortex. Therefore, the discriminative performance of certain descriptor is sufficient in separating neurons in different cortical areas of the rat brain, but more importantly, is powerful enough in revealing sub-clustering in a population of neurons.

### Morphological aberrations

Revealing morphological aberrations resulting from neurodevelopmental and acquired disorders is an important step in understanding the pathophysiology of these diseases. Thus, unbiased methods to distinguish and separate normal neuronal morphology from aberrant morphology becomes essential. For that purpose, **Dataset 5** included pyramidal neurons in layer 5 of rat somatosensory cortex in control (n=20) and experimental condition (n=16) was assembled. This study assessed morphological changes of cortical pyramidal neurons in hepatic encephalopathy. Interestingly, the authors report that although dendritic arbors remained unchanged in rats with hepatic encephalopathy, dendritic spine density was significantly reduced [30]. Indeed, as one would expect, the detection rates based on all the descriptors was low (Table 1), suggesting that neurons from the control group and the experimental group were virtually indistinguishable. Unsurprisingly, the combined descriptor approach (Supplementary Fig. 7b) (see Supplementary §4.2) resulted in intermingling of neurons from the control and experimental group. These results confirm previous findings that neuronal morphology is largely unaltered in rat cortical neurons with hepatic encephalopathy. More importantly, although the study only assessed path length and number of terminal ends in control versus experimental condition, we reveal using multiple Sholl descriptors that additional parameters related to neuronal morphology are in fact comparable between the two neuron groups. Nevertheless, although our descriptors do not reveal any structural differences between neurons in the two groups, there may be other features that differ which our descriptors do not capture. In a future study, we intend to construct additional Sholl descriptors that may potentially reveal structural differences between control and experimental neurons in this dataset.

### Classification and metric learning

Finally we apply our descriptors to a relatively large dataset of 359 neurons in order to classify them. **Dataset 6** is comprised of: hippocampal granule (n=77), hippocampal microglia (n=79), spiny (n=66), purkinje (n=60), and interneuron (n=77). Ametric learning classification scheme is then used to generate a suitable metric that can differentiate among the classes. This is accomplished by first applying all Sholl descriptors on this dataset to determine which ones result in the best detection rates (‘feature selection’). We subsequently choose these features (i.e. descriptors) for inclusion in our classification scheme. Specifically, Table 1 shows that the energy descriptor was ineffective in distinguishing among classes in this dataset, as evidenced by low detection rates (below 80%). Therefore, this descriptor is removed, and only the descriptors that perform well in separating these classes are included (flux, wiring, leaf index, branching pattern, tortuosity and TMD). Next, we vectorize the neurons (Supplementary §8) based on these descriptors, resulting in classes of vectors in Euclidean space (in ℝ^30^). We then use a t-SNE plot to visualize our high dimensional descriptor vectors in 2D space (Fig. 6a). Some neurons from different classes in this dataset are clearly overlapping and poorly separated. Data is then fitted and transformed into a new metric space using the Large Margin Nearest Neighbor (LMNN) metric learning algorithm which learns a Mahalanobis distance metric in the K-Nearest Neighbor (KNN) classification setting [31] (see Supplementary §4.3). In order to cross validate the model, 70% of data is randomly selected for the training phase and the rest is used for test. A model accuracy score of 95% is obtained. In order to test for over-fitting we use a k-fold cross validation technique where the training set is split into k equal sets or folds. Each k fold is used as a validation set and the k-1 remaining folds are used for training. We used a 10-fold validation method which resulted in an average performance score of 92%. This powerful new approach results in excellent separation of classes in this dataset. Fig. 6b shows the data plotted in the new transformed space.

**Fig. 6.**
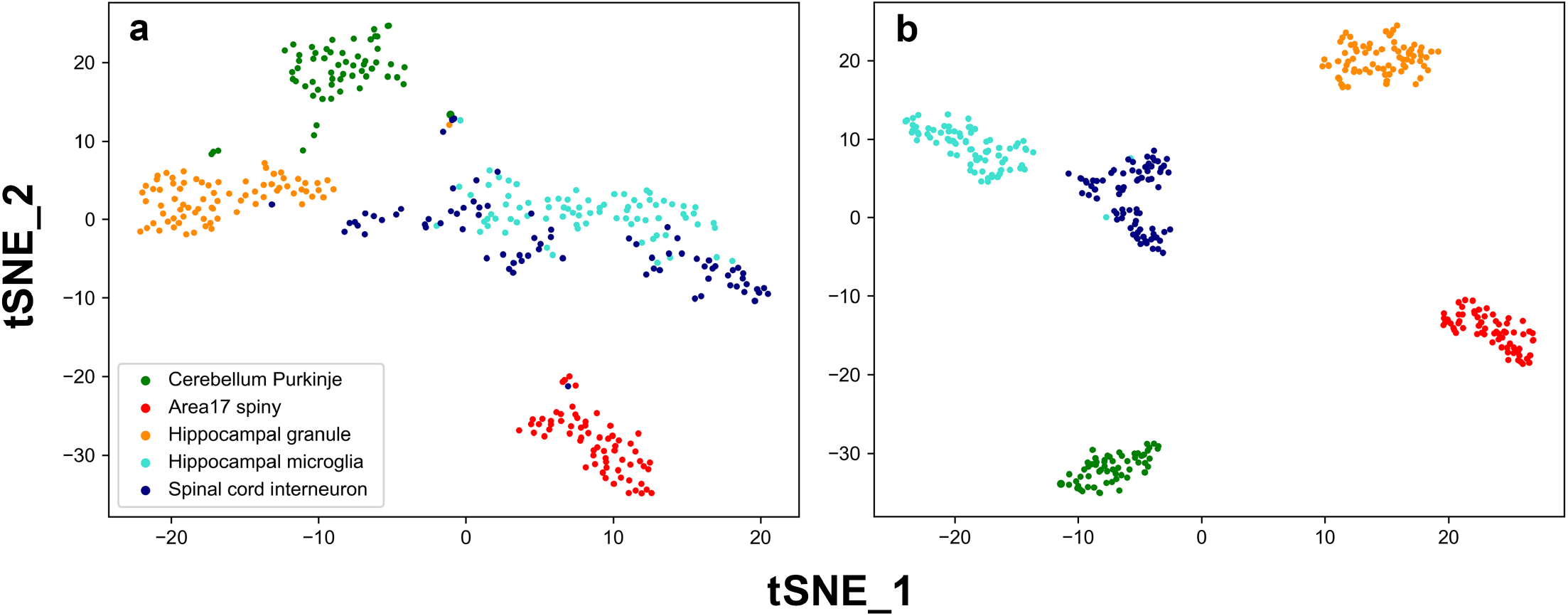
Classification and Metric learning for Dataset 6. Each dot represents a neuron which is color coded according to its class. tSNE was performed to reduce dimensions. (**a**) Vectorized neurons in Euclidean space. (**b**) KNN classification in the new metric space.

## 4. Discussion

In this work we introduced novel descriptor functions of tree structures and applied them for classification of neuronal cell types. Importantly, we obtained substantially better clustering results when we compared the performance of our descriptors with conventional methods. The essence of our methods is, for every neuron, to construct a function having values in some metric space (Sholl function). This Sholl function captures the evolution of a particular morphological feature as distance from the soma increases. Sholl functions of different neurons can be compared and the distance between them informs of how (dis)similar given neurons are with respect to the morphological featured captured by the applied Sholl functions. We illustrate that certain descriptor functions can effectively cluster classes of neurons with subtle morphological variations, as well as discriminate among widely different classes of neurons in agreement with expert assignment. Additionally, we leverage metric learning techniques to provide more robust classification. Our framework is powerful enough to separate diverse classes of neurons across different brain regions and species. Our results reveal several key findings regarding the considered datasets.

The six representative datasets used in this study, derived from different areas, layers, and species, were chosen to ensure morphological diversity and prove wide applicability of the presented techniques. In Dataset 2 (different types of neurons taken from different regions of the mouse brain), we show that the TMD and tortuosity descriptors performed very well in recovering the clusters corresponding to the brain regions the neruons came from. Specifically, based on the tortuosity descriptor the Martinotti and pyramidal cells each formed their own cluster. Interestingly, dendritic tortuosity has been shown to vary among different non-pyramidal neuron classes in the rat brain, whereby Martinotti cells in layer II/II and V of the frontal cortex have higher tortuosity than other cell types [32]. In the mouse brain, dendritic tortuosity increases as a function of increasing branch order on apical dendrites of hippocampal CA1 pyramidal cells [33]. Additionally, dendritic tortuosity of layer II/III pyramidal cells appears to increase from caudal to rostral regions in mouse cortex [34]. Our tortuosity descriptor is therefore robust in detecting subtle differences in dendritic tortuosity among neuron groups. The upper limit for tortuosity values appears to be 2, which is consistent with prior reports [32]. Importantly, we improved discrimination accuracy by using a combination of descriptors which effectively assigns weights to the function with the best separation results.

Anatomical studies have shown that interneuron morphology is highly diverse in the cerebral cortex. For example, interneurons with similar somatodendritic morphology may differ in axonal arborization patterns [13]. Therefore, axonal morphometric features are typically required for accurate classification of interneurons as they have been shown to capture important differences among interneuron subtypes [35]. We did not analyze axonal features in our descriptors as that could explain why Martinotti, and medium spiny neurons were largely intermingled (Dataset 2). However, based on dendritic features alone some of our descriptors (tortuosity and TMD) were able to reliably distinguish interneuron subtype (Martinotti) from other neuronal cells types such as purkinje and retinal ganglion cells (Dataset 2). In a future study, we will focus our efforts on interneuron subtypes in order to incorporate important axonal features into our descriptors for a more accurate classification scheme.

Prior work has revealed regional differences in pyramidal cell morphology in the monkey brain. Specifically, in the Old World macaque monkey, pyramidal cells become progressively larger and more branched with rostral progression through V1, the secondary visual area (V2), the fourth visual area (V4), and inferotemporal cortex (IT) [4, 8, 37]. Therefore, we were interested in testing whether our descriptors can detect differences in the morphology of pyramidal cells from different visual cortical areas of the vervet monkey (Dataset 3), another species of Old World monkeys. Indeed, we find that the performance of the branching pattern descriptor results in excellent clustering of cells from V1, V2, and V4. This suggests distinct differences in the branching pattern of basal dendrites of pyramidal cells residing in these areas. The wiring descriptor which is a proxy for total dendritic length yielded reasonable clustering of neurons, with some intermingling of neurons from all areas. This is not surprising given that in some species such as the tree shrew, differences in pyramidal cell morphology throughout the visual cortical hierarchy is less pronounced [38]. Even in rodents, regional differences in pyramidal cell morphology appear to be less noticeable than in primates [39, 40]. Therefore, the fact that pyramidal cells in V2 are intermingled with cells in V4 in our cluster analysis based on the wiring descriptor reflects genuine similarities between these two population of cells, and further suggests inter-neuron variations within each visual cortical area.

Collectively, the results from this study highlight the robustness of our framework in quantitatively characterizing and discriminating among different neuronal cell types. Certain morphological features and thus specific descriptors are better suited in separating distinct neuronal cell types. For instance, the branching pattern descriptor appears to perform very well in detecting most neuronal cell types. This descriptor measures how far or how fast nodes appear (bifurcations) and disappear (leaves), as measured from the soma. Conversely, the energy descriptor, which reveals the distribution of nodes around the soma, appears to reliably detect retinal ganglion cells, purkinje, and interneurons. Importantly, our use of metric learning techniques resulted in more optimal classification (Fig.6). Progress in the development of unbiased clustering methods to distinguish among groups of neurons will further our understanding of the relationship between brain structure and function. The toolkit of morphological descriptors introduced here, and the development of new methods will potentially lead to the discovery of novel sub-classes of neurons [41]. Additionally, our descriptors will aid efforts to uncover differences between normal and aberrant neuron morphology which is commonly associated with various disease states. For instance, changes in dendritic morphology have previously been described in a number of disease states, including Alzheimer’s disease [15], schizophrenia [42], and mental retardation [43]. Given that our tool kit of descriptors discriminated among different types of cells as well as revealing subclasses of cells, its utility may be extended to the study of brain diseases potentially identifying which subtypes may be affected in various disease states.

## 5. Software

MATLAB v2019a (The Mathworks Inc., Natick,MA. RRID: SCR_001622), Python v3.7 (Python Software Foundation. RRID:SCR_008394) and R v4.0.3 (R Foundation for Statistical Computing. RRID:SCR_010279) were used for computations

## 6. Code availability

Code for the data analysis was deposited to the Github repository and is available at https://github.com/reemkhalilneurolab/morphology

## 8. Acknowledgements

We thank Mohammad Rowaizak for his help with figure 1,2 and 3 illustrations.

## 10. Author contributions

S.K, R.K. and A.F. conceptualized the project; S.K, A.F., and P.D designed the computational analysis; P.D. and A.F. performed all the coding. S.K, R.K., P.D and A.F. interpreted the results; R.K. and S.K wrote the initial draft paper; S.K, R.K., P.D and A.F. revised the initial draft and wrote the final paper.

## Supplementary Material

## 1. Summary of Methods Used to Classify and Cluster Neurons

Sholl descriptors form a toolkit to analyze neuronal morphology. Descriptors combined with standard hierarchical clustering methods, a detection algorithm (also used for feature selection), grid search, and metric learning functions are used to cluster and classify a dataset of neurons.

1. Analysis based on a single descriptor:
  - (Clustering) Given a dataset of unlabeled neurons we can use a particular descriptor in order to cluster them according to that descriptor. For example, if *ϕ* = *T* is tortuosity, we can set the distance matrix for the associated Sholl metric *d*_*T*_ and then run standard hierarchical clustering to obtain dendograms. The obtained dendrogram reveals whether neuronal cell types differ according to their tortuosity (i.e. cluster together), or if their tortuosity is comparable (cells from different neuron types will be intermingled).
  - (Detection) It is a simple distance-based measure designed for a purpose of this paper. It assess the performance of a given descriptor in identifying neurons with a given label within a dataset. Its idea is to find a ball *B* that contains at lest *p*% of all neurons with the label *l* and, in addition, the neurons of the label *l* compose at least *p*% of all neurons in *B*. See Figure 1 for illustration and the precise construction in §1.4). Detection rates can be used as feature selection when running classification schemes.
2. Analysis based on a combination of descriptors (details in §4):
  - (Vectorization and unsupervized clustering) A given neuron can be converted into a vector using our descriptor functions and metrics. There are different approaches that may be used for that purpose We have chosen to use an approach detained in §4.1. Given the feature vectors, we then run a standard clustering algorithm.
  - (Classification) Given a dataset of neurons distributed among a number of classes, and a number of morphological features, we can determine which class a newly introduced neuron is associated with (i.e. shares the most features with). More precisely, suppose we are given classes of neurons *C*_*i*_, 1 ≤ *i* ≤ *n*, and morphological descriptors *ϕ*_1_, …, *ϕ*_*k*_ which can be measured for all neurons. Therefore, given a newly introduced neuron *N*, we can determine which class, with measurable likelihood, this neuron belongs to.
  - (Differentiation) By linearly combining descriptor metrics, we can separate classes and obtain a new clustering metric. This approach is similar to metric learning in that it will separate different classes, unless the classes are truly indistinguishable.

### 1.1. Normalization

A given Sholl descriptor 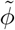 can be normalized so that it is supported on [0, 1]. More precisely, let 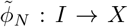 be the descriptor function for a neuron *N*, where *I* = [0, *R*(*N*)] or [0, *L*(*N*)]. If 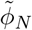 is supported on [0, *R*(*N*)] for example, where *R*(*N*) is the span of the neuron (see §2.2), we define the corresponding normalized descriptor *ϕ*_*N*_ by:

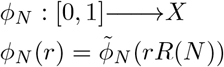

If 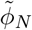 is a function of path length, and thus supported on [0, *L*(*N*)], then we normalize in the same way, with *L*(*N*) replacing *R*(*N*). If 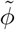 is isometry invariant to begin with, then so is *ϕ*. Being isometry invariant means that for any linear transformation *A* of ℝ^3^, *ϕ*_*AN*_ (*r*) = *ϕ*_*N*_ (*r*), where *AN* is the image of *N* ⊂ ℝ^3^ by the transformation *A*.

All our descriptors are normalized and supported on [0, 1]. The constructions we provide are such that all real-valued Sholl functions we consider are step functions. Fig. 4 illustrates these step functions for a very simple granule cell in mouse olfactory bulb.

### 1.2. Functional Metrics

Each Sholl descriptor defines a pseudo-metric on any given set of neurons 𝒩. Let *ϕ* be a normalized Sholl descriptor which associates to each neuron *N* a descriptor function *ϕ*_*N*_ : *I* →*X, I* = [0, 1], and *X* a metric space. Every Sholl descriptor *ϕ* gives rise to a map

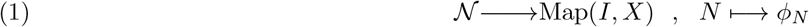

Let *d* be any metric on the space of functions Map(*I, X*). It induces a pseudo-metric *d*_*ϕ*_ on 𝒩 by setting

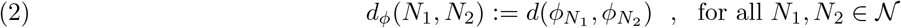

Note that, since *ϕ* is *O*(3)-equivariant, we have that *d*_*ϕ*_(*N*_1_, *N*_2_) = 0 if *N*_1_ is the image of *N*_2_ by a rotation or a reflection. It is possible to upgrade *d*_*ϕ*_ to a genuine metric on the set of neurons up to isometry, under the assumption that (1) is an injective map. This is true in practice. This fact is important since it allows us to conclude the following: *The smaller is d*_*ϕ*_(*N*_1_, *N*_2_), *the more similar are N*_1_ *and N*_2_ *in the feature ϕ*. This statement is at the basis of our stability results (see Supplementary material §2).

The functional distance *d* on the space Map(*I, X*) we choose to work with is the “*L*^1^ distance”

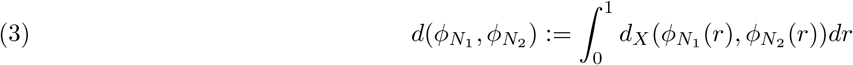

All *ϕ*_*N*_ constructed in this paper are real-valued *step functions*, except for the Sholl-TMD in §1.6.8. We give the formula for the distance in this case. Let *ϕ*_1_, *ϕ*_2_ be two step functions with jumps at radii *r*_1_, …, *r*_*q*_ and *s*_1_, …, *s* respectively. This means that *ϕ*_1_ is constant on [*r*_*i*_, *r*_*i*+1_[, and similarly *ϕ*_2_ is constant on [*s*_*j*_, *s*_*j*+1_[. Let

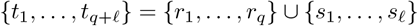

and order the *t*_*i*_’s by increasing magnitude so we can assume

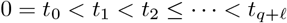

Then the *L*^1^-distance between the step functions is given by

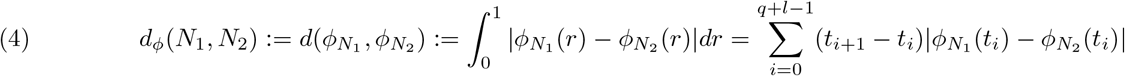

Other choices of metrics we can work with for real valued Sholl functions are the *L*^*p*^ metrics for *p >* 1 or the Sup metric 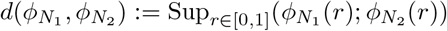. Once we can measure functional distances between descriptor functions, we can measure “Sholl distances” between neurons as indicated in (2). These distances are then used to cluster and classify neurons.

### 1.3. Clustering

The ultimate goal is to find measurable and quantifiable morphological differences between classes of neurons. When given a Sholl descriptor, and a selection of neurons *N*_1_, …, *N*_*q*_, the standard procedure is to generate a distance matrix associated to the descriptor

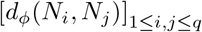

This is a symmetric matrix with non-negative entries, and zeros along the diagonal. Any such distance matrix produces a dendrogram using standard hierarchical clustering algorithms. It is not reasonable to expect a single descriptor to cluster faithfully a given set of classes of neurons (Supplementary Fig. 4). The advantage of developing multiple descriptors based on various morphological features reveals which features are uniform and which are different among classes of neurons. Combining descriptors to differentiate between classes of neurons is another method we use. This combination can be achieved at the level of distance matrices, or at the level of Sholl descriptors since these form a vector space (in fact an algebra) of functions. Indeed, given two normalized Sholl descriptor functions *ϕ*_1_, *ϕ*_2_ : *I* →ℝ, we can take linear combinations. This sum is also stable, as defined in §2, if we start with stable descriptors.

### 1.4. Detection and Feature Selection

Let *C*_1_, …, *C*_*k*_ be *k* distinct classes of neurons. Neurons of each class can be compared with neurons from other classes. We say that a descriptor *ϕ* has at least an *n*% level of detection of a class *C*_*i*_ if there is a ball *B*^*ϕ*^ in the *d*_*ϕ*_ metric so that more than *n*% of all elements of *C*_*i*_ are within *B*^*ϕ*^, and of all elements in *B*^*ϕ*^, more than *n*% are from *C*_*i*_.

#### Example 1.1.

Suppose we have three classes of neurons *C*_1_, *C*_2_, *C*_3_, each consisting of 5, 4 and 3 neurons respectively. Let *ϕ* be a given Sholl descriptor, and suppose there is an *ϵ* ball in the *d*_*ϕ*_-metric, that contains 4 elements of *C*_1_ and 2 element from *C*_2_ ∪ *C*_3_. This ball contains 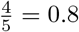 of the total of all *C*_1_-neurons (i.e. 80%), while 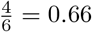 or 66% of all neurons in this ball are *C*_1_-neurons. We say that the descriptor *ϕ* has detected *C*_1_ to a 66% level at least, which is the lower percentage from among 80% and 66%. Figure 1 illustrates this construction on these three classes.

The detection algorithm is described in §3. We also use detection as a method for feature selection when we run several descriptors on a given set of classes. The set of descriptors with detection rates that are less than a certain percentage are deemed ineffective in differentiating among these classes and can thus be excluded from further analysis (see §4.3).

### 1.5. Combination of Descriptors and Classification

A single descriptor may detect features within a family of neurons, but it alone may not be able to differentiate between many classes at once. The idea of “combining” several descriptors together into one single descriptor offers a more effective tool in differentiating between classes. The Sholl descriptor metrics are perfectly well-suited to provide such a classification. We have devised three combination methods, each being applicable within its own specific context. This was discussed at the beginning of §3 and the details can be found in the supplementary material §4.

**Fig. 1.**
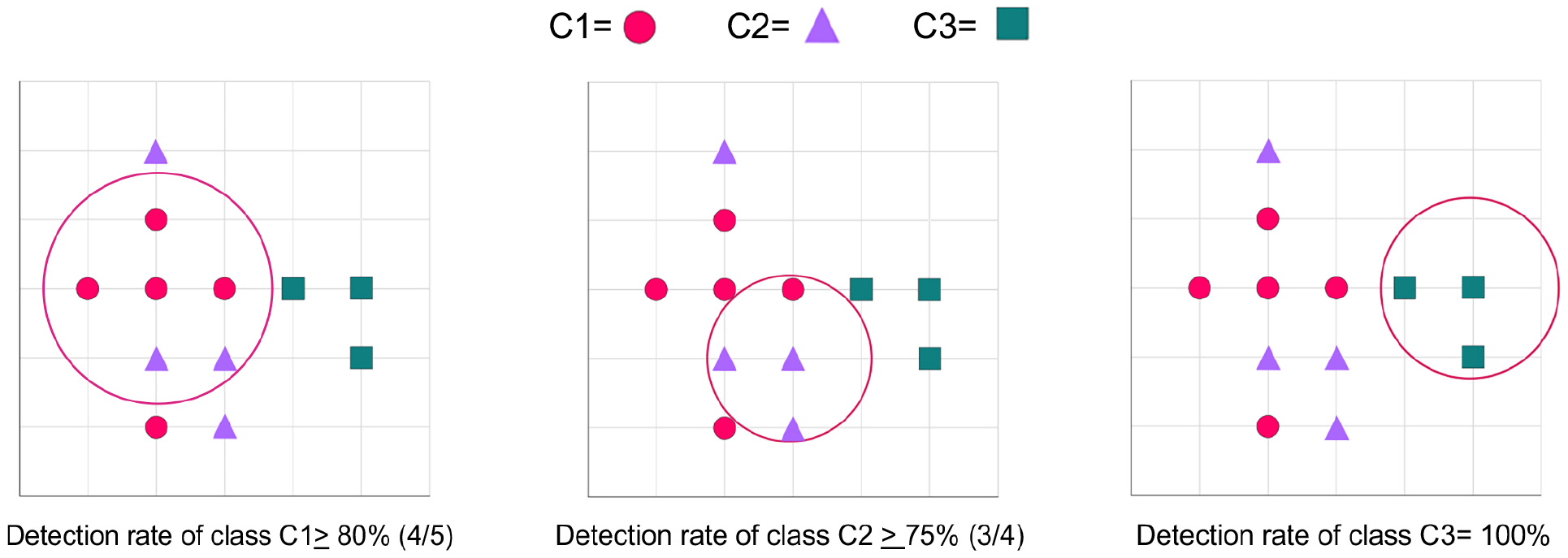
Method used to determine detection rate. Each circle is the boundary of a disk in the Euclidean metric.

### 1.6. The Sholl Descriptors

Given a neuron *N* viewed as an embedded tree in space, we define

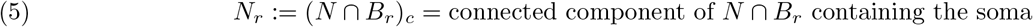

Here *B*_*r*_ is the ball of radius *r* around the soma.

#### 1.6.1. The Branching Pattern Descriptor

This morphological descriptor detects patterns that results from the distribution of branches and leaves relative to the soma (see Fig. 2).

Let *N* be a neuron which we view as a collection of single rooted binary trees in ℝ^3^, with the common root being the Soma. Label *B*_1_, …, *B*_*q*_ the branch points of *N* and label all leaves by *L*_1_, …, *L*_*k*_. Let *r >* 0 be the radial distance measured away from the Soma. Order the branch points and leaves by increasing *r*, so that if *r*_*i*_ indicates the distance of the *i*-th node to the soma, we have 0 *< r*_1_ *< r*_2_ *<… < r*_*k*_ (equal radii can be removed by an infinitesimal perturbation).

Fixing a neuron *N* as before, associate to each *r* ∈ *I* the number *α*(*r*) defined by

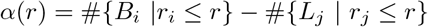

Let *R*(*N*) be the span of the neuron *N* and define the function

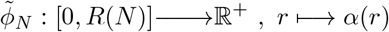

This is a step function since *α*(*r*) is constant on intervals *r*_*i*_ ≤ *r < r*_*i*+1_. The normalized version takes the form

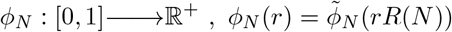

This defines our branching-pattern descriptor *ϕ*. This descriptor is isometry invariant (it only depends on the distance of the branchpoints from the soma) and so it is well-defined as a function on the isometry classes of neurons as already indicated. Note that

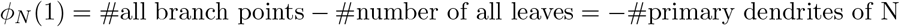

since the number of primary branches is the difference between the number of leaves and the number of bifurcation points.

##### Example 1.2.

In Fig. 2, Tree structures of two different neurons are chosen (A) pyramidal and (B) stellate. The corresponding Sholl descriptor functions reveal obvious difference (C). The red curve which depicts the branching pattern of the pyramidal cell reveals that branching occurs rapidly close to the soma, but much slower as you move further away from the soma. Conversely, branching for the stellate cell is changing uniformly and steadily as you move away from the soma. Both neurons have similar branching counts: neuron (A) has 21 bifurcations and 30 leaves, while neuron (C) has 32 bifurcations and 49 leaves. This gives that 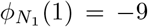 and 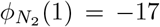 as depicted.

#### 1.6.2. Tortuosity Descriptor

Given a representation of a neuron by an embedded tree in space, let us label its nodes by *P*_1_, …, *P*_*n*_ ∈ ℝ^3^. For any two nodes, we can consider both path distance and Euclidean distance between them.

**Fig. 2.**
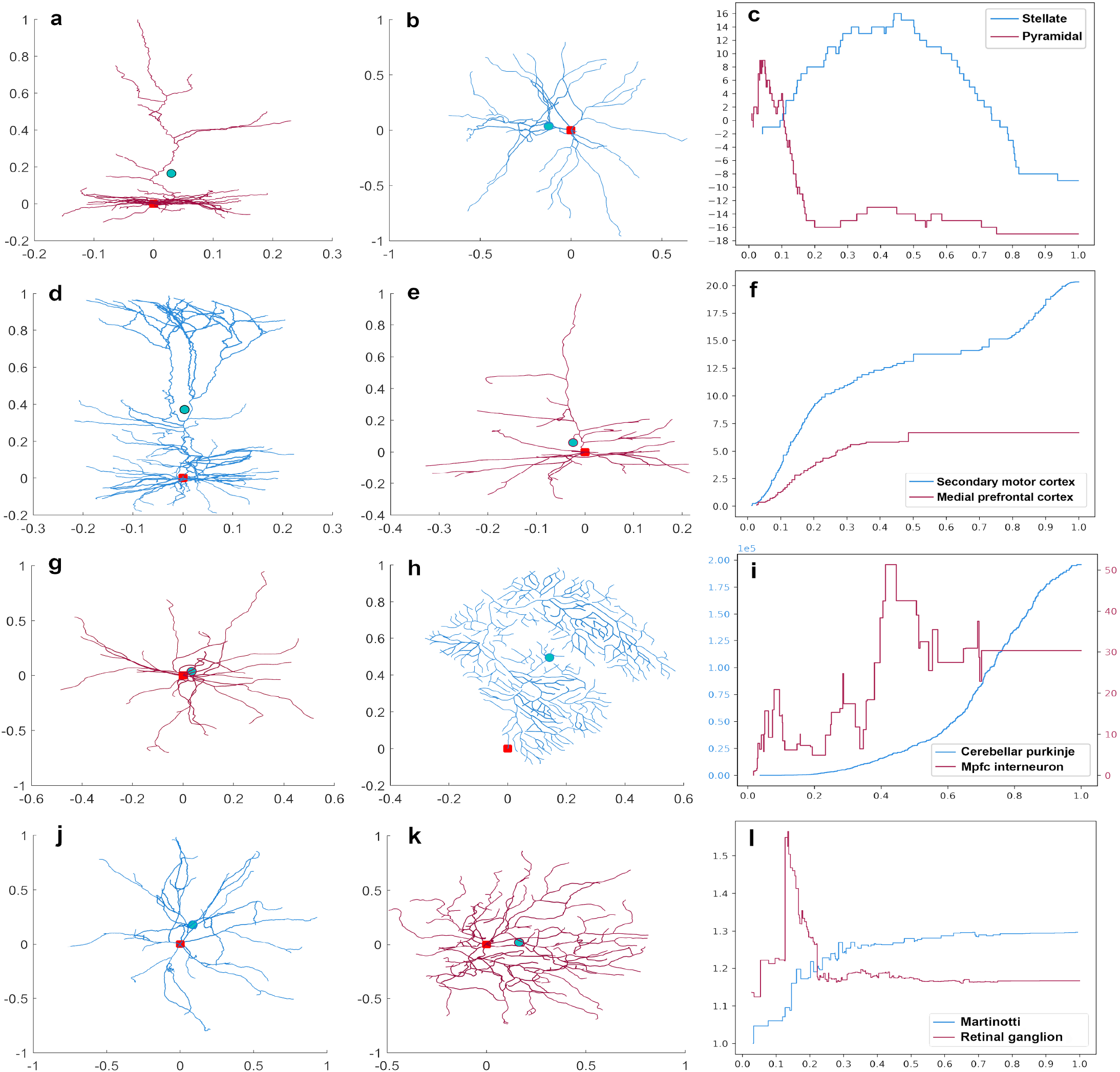
Representative neurons and the corresponding step functions. (**a**) Pyramidal and (**b**) stellate cell. (**c**) The step functions for each neuron generated from the branching pattern Sholl descriptor. The branching Sholl functions show that neuron (**a**) is branching quickly near the soma, and leaves appear much closer to the soma than for neuron (**b**). The number of primary branches for each neuron is the value of the corresponding function at 1. (**d**) Pyramidal cell in secondary motor cortex and (**e**) in mPFC. (**f**) The step functions for neurons (**e**) and (**d**) generated from the wiring Sholl descriptor. The sharp increase towards the end for the wiring Sholl function for cell **d**. reveals the existence of an apical tuft. (**g**) Interneuron and (**h**) Purkinje cell. (**i**) The step functions for each neuron generated from the energy Sholl descriptor. The two energy Sholl functions show completely distinct features. We obtain maximal energy values for purkinje cells (a defining feature). The total energy value at the soma for neuron (**g**) is 200000 units, while this value is 50 for neuron (**h**). (**j**) Martinotti and (**k**) Retinal ganglion cell. (**i**) The step functions for each neuron generated from the tortuosity Sholl descriptor. The red dot represents the soma and the green dot is the barycenter of all nodes

If *P*_*i*_ is a parent node and *P*_*j*_ is the child node, let *b*_*i,j*_ be the dendritic path distance between these nodes, and let *d*_*i,j*_ be the length of the segment [*P*_*i*_, *P*_*j*_]. The ratio of both distances is

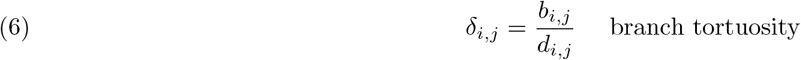

Consider a neuron *N* with *n* nodes §2.2. Let (parent, child) be a pair of adjacent nodes. There are exactly *n* − 1 such pairs coinciding with the number of branch segments. Define the average tortuosity of *N* to be the average

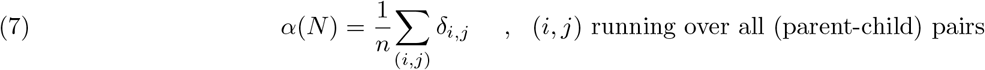

It is clear that 1 ≤*α*(*N*) for all choices of *N*. The (non-normalized) Sholl descriptor function associated to this construction is now given as follows: order the the nodes of *N* by increasing radii as before. Then define

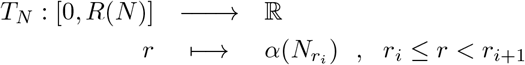

where *B*(*r*) is the ball of radius *r* around the soma. We then take the normalized version §1.1.

#### 1.6.3. Taper Rate Descriptor

We start with a neuron *N* and list all *path distances* of the nodes to the soma in increasing order 0 *< ℓ*_1_ *< … < ℓ*_*k*_. Each node has a dendritic thickness (or width) that tapers as we move away from the soma along the dendrite. We can measure the tapering rate as a function of path distance. More precisely, define

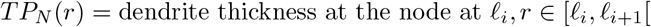

and then take the associated normalized Sholl descriptor by dividing *ℓ*_*i*_ by the length of the longest dendrite. This is a Sholl function whose variable is path length and not radial distance.

#### 1.6.4. Flux Descriptor

We define the Sholl descriptor *F* and the associated flux functions *F*_*N*_ : [0, 1] →ℝ for a given neuron *N*. Let 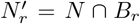, where *B*_*r*_ is a ball of radius *r* centered at the soma. Notice that 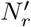 can be different from *N*_*r*_ (5) if there are dendrites that leave *B*_*r*_ and then enter again. If a dendrite crosses the boundary sphere *S*_*r*_ at a point *P* ∈ *N* ∩ *S*_*r*_, we identify the parent of *N* (inside the sphere) and the child of *N* (outside the sphere). So the parent and child are on either sides of the sphere. The direction vector 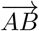 from father to child points outward if *A* is inside, and points inward if *A* is outside the sphere. Consider the segment [*A, B*] and let *C* be the point on the segment that cuts the sphere. We then assign the value

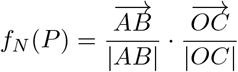

This is the cosine of the angle between the unit vector along 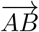 and the normal to the sphere going through *C*. This value is maximal if 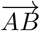 is aligned with the radial vector at *C* and the angle is zero.

To define the total flux function, order the nodes of *N* as before by increasing values of their distances from the soma 0 *< r <* … *< r*. For every 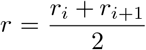, take the sphere of that radius *r*, look at all dendrites intersecting that sphere at *P*_1_, …, *P*_*k*_ and add up the values obtained from the construction outlined above. This value is

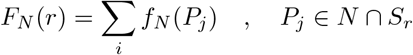

This gives rise again to a step function *F*_*N*_ : [0, 1] → ℝ by setting

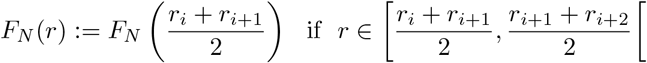

Additionally, if *A* = (*a*_1_, *a*_2_, *a*_3_) is the parent marker and *B* = (*b*_1_, *b*_2_, *b*_3_) the child marker, such that either |*OA*| *< r <* |*OB*| or |*OB*| *< r <* |*OA*|, that is on different sides of the sphere, the point of intersection *C* = (*c*_1_, *c*_2_, *c*_3_) of that sphere with the segment [*A, B*] is obtained by setting 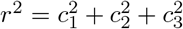 and *C* = (1 − *t*)*A* + *tB*, then solving for *t* through a quadratic. The flux value at *C* is

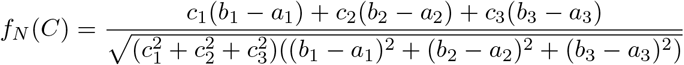

#### 1.6.5. The Leaf Index Descriptor

From each node grows a new dendritic tree with a number of terminal points. Counting the number of these terminal points for each node *P* gives the “leaf index” of *P*, and we write it as li(*P*). When *P* is a leaf, we set li(*P*) = 1.

As before, given a neuron *N*, order its nodes *P*_1_, …, *P*_*q*_ by increasing order of path distance to the soma 0 *< ℓ*_1_ *<* … *< ℓ*_*k*_, and define the Leaf Index Sholl Descriptor as follows:

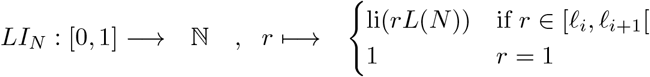

Evidently *LI*_*N*_ (1) = 1, which is the value at the furthest leaf, while *LI*_*N*_ (0) is the total number of leaves. This is again a step function and distances between leaf index Sholl functions can be given by the standard formula (4).

#### 1.6.6. Total Wiring Descriptor

“Total wiring” is a morphological feature which measures the total dendritic length of neurons. When used as a Sholl descriptor, it gives total length of dendrites, but also their density as we move away from the soma.

Given a neuron *N*, let *tℓ* (*N*) be the total length of all dendrites of *N*. If *N*_*r*_ is part of the neuron within a sphere of radius *r* from the soma (5), then

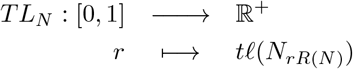

This is a normalized Sholl function, which always starts at value 0 and ends up at value *TL*_*N*_ (1) = *tℓ* (*N*) which is the total wiring of the neuron. As for other Sholl functions, we will only consider the step function version of this construction, where once more one defines for *r* ∈ [0, 1],

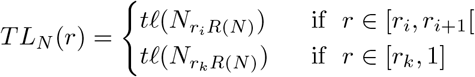

where 0 *< r*_1_ *<* … *< r*_*k*_ ≤ 1 are the normalized radial distances of the nodes listed in increasing order.

#### 1.6.7. Energy Descriptor (Nodal Distribution)

Given a neuron *N*, consider all its nodes as cloud points in 3D space viewed as charged particles. The charge each node carries will be proportional to the thickness of the branch at that point, if available and 1 otherwise. These charged nodes affect the space around them through the electric field they generate. This electric field is a well-defined map

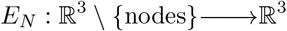

Taking the intensity of the vector field at each point of ℝ^3^ \{nodes} gives us a measure of how space is being affected by the neuron. This also gives a measure of how the nodes are distributed in space as we will later illustrate in the case of Purkinje cells.

Let *ζ* := {*P*_1_, …, *P*_*n*_} be the nodes of *N*, with *P*_*i*_ having charge *q*_*i*_. This charge *q*_*i*_ is chosen to be the width of the dendrite at point *p*_*i*_. Each point *P*_*i*_(*x*_*i*_, *y*_*i*_, *z*_*i*_) of *ζ* contributes an electric vector field which is normalized to have length *q*_*i*_ and which is of the form

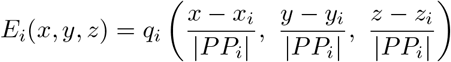

where 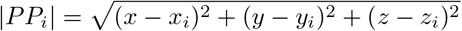. By superposition, the node configuration *N* gives rise to a vector field *E*_*N*_ (*x, y, z*) =∑_*i*_ *q*_*i*_*F*_*i*_(*x, y, z*) with square intensity

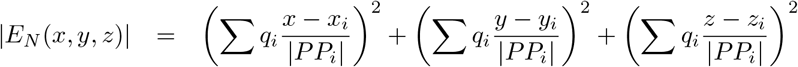

Let *O*(2) be the group of orthogonal matrices. This group acts on ℝ^3^, and thus on the set of neurons. If *A* ∈ *O*(2) and *N* ⊂ ℝ^3^ a neuron, we write *A*(*N*) the image of *N* under this action.

##### Lemma 1.3.

*The intensity at the soma*, |*E*_*N*_ (0, 0, 0)|, *is* 0(2)*-invariant*.

*Proof*. We show first that |*E*_*A*(*N*)_(*P*)| = |*E*_*N*_ (*A*^−1^(*P*))| for *P* ∈ ℝ^3^. The nodes of *N* are {*P*_1_, …, *P*_*n*_}. Write 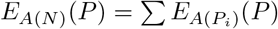. Since 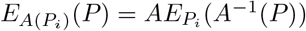, and since *A* is linear and preserves lengths, it follows that

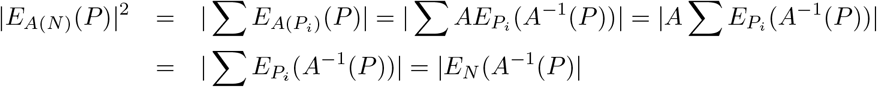

At the soma *P* = (0, 0, 0), *A*^−1^(*P*) = *P*, so that |*E*_*A*(*N*)_(*P*)| = |*E*_*N*_ (*P*)|, which is what is claimed. □

**Fig. 3.**
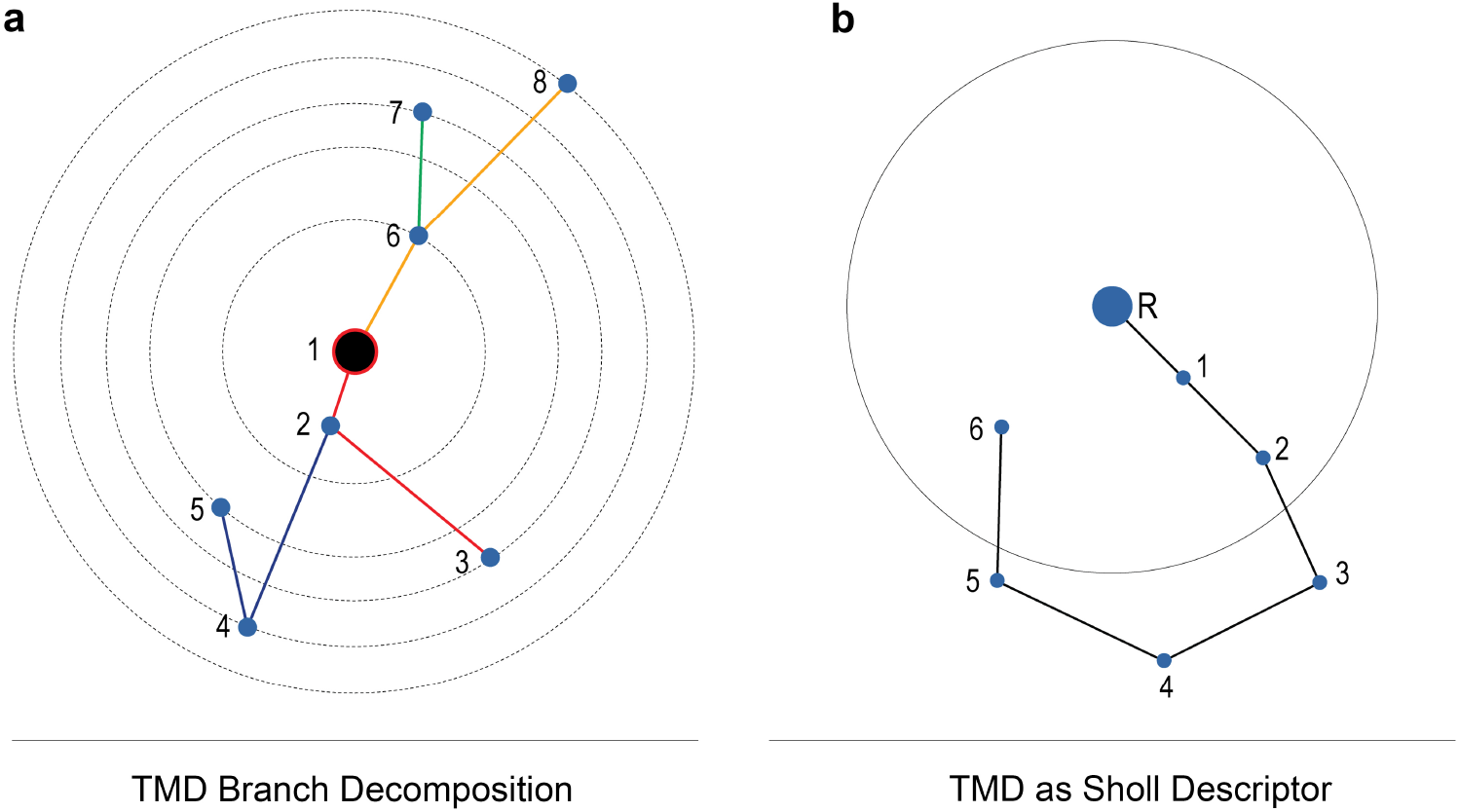
(**a**) Example of TMD-path decomposition on a simple planar tree. The soma marked with 1 is the root. Equicentered circles reveal the distances of nodes from the root. The furthest node is node 8. The paths from the TMD-path decomposition are: {[5, 4, 2], [3, 2, 1], [8, 6, 1], [7, 6]} (**b**), The tree *T* with a single path *x* starting at the root *R*. When using TMD as a Sholl-type descriptor by considering TMD of *T* ∩ *B*(*R, r*) we will only see the final barcode [0, *d*(*R*, 6)] for *r* ≥ *d*(*R*, 4). For the radii *r* between *d*(*R*, 6) and *d*(*R*, 4) the endpoint of the persistence interval will be equal *r*. When *r* reaches *d*(*R*, 4) the endpoint of the persistence interval it will then jump down to *d*(*R*, 6).

Our Sholl descriptor associates to every neuron the map

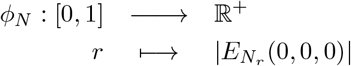

where *N*_*r*_ is as in (5). This map adds the unit vectors at the soma, one for each node, and takes the magnitude. We can also think of energy as the effect of the nodal distribution around the soma. If all nodes are on one side of a plane going through the soma, then their contributions is greatest (eg. Purkinje cells have very large energy values), as opposed to nodes that are evenly distributed around the soma. In this latter case, several cancellations occur and the energy value tends to be small.

#### 1.6.8. The Topological Morphological Descriptor (TMD)

Let *T* ⊂ ℝ^3^ be a tree with a root *R*. A *path* is any continuous sequence of edges in *T*. Each path *x* have unique initial *b*(*x*) and terminal *d*(*x*) vertices. The TMD is based on a method of decomposing a given tree *T* into a collection of paths such that the sum of those paths is the whole tree *T*. In addition, any two paths from that decomposition will either have empty an intersection, or their intersection is the endpoint of one of them (and in this case a branchpoint of *T*).

The TMD path decomposition is obtained using the following procedure; All the paths from the TMD-path decomposition starts at the leaves of *T*. They continue along the tree, towards the root, until they reach a node *n* of degree 3 or higher in *T*. In the node *n* all the paths except from one terminate. The path that continues through the node *n* is the one with the initial node further away from *R* (soma)^2^. Once a path reaches the root *R*, it does not continue any further (it terminates). For example, a TMD-path decomposition is presented in Figure 3a.

Given a TMD-path decomposition as described above, a collection of pairs of numbers inspired by *persistent diagram* is associated to this decomposition. For that purpose, a path *x* having the initial and terminal vertices *b*(*x*) and *d*(*x*) correspond to persistence interval [*d*(*R, b*(*x*)), *d*(*R, d*(*x*))]. Using the terminology from persistent homology, we say that the path *x* is *born* at the radius *d*(*R, b*(*x*)) and *dies* at the radius *d*(*R, d*(*x*)). The collection of all such birth-death pairs is then used as a signature of the tree *T*. As the obtained signature has the structure of persistent diagram, we further adopt various metrics from persistent homology to compare such diagrams.

To fit the TMD into the scheme of the current paper we will now turn it into a Sholl descriptor having values in the space of persistence diagrams. For that purpose let *T*_*r*_ be the connected component of *T* ∩*B*(*R, r*) containing *R*. Let us make two simple observations:

1. Note that the TMD-path decomposition of *T* restricted to *T*_*r*_ is a valid TMD-path decomposition of *T*_*r*_. To see that, let us consider a branching node *n* in *T*, such that *d*(*R, n*) ≤ *r*, and the same node in *T*_*r*_. The paths from *n* to the leaves in *T*_*r*_ will either be the same as in *T*, or they will be cut short in *T*_*r*_ by *B*(*R, r*). In both cases, the path that does not terminate at *n* in *T* will also be the path with the initial point further away from *R* in *T*_*r*_. It is possible that more than one path in *T*_*r*_ joining in *n* will be cut short by *B*(*R, r*). However in this case we can choose to continue in *T*_*r*_ the same path that continues in *T*. Consequently, a TMD-path decomposition in *T*_*r*_ can be obtained by appropriate restriction of the TMD-path decomposition in *T*.
2. Suppose we consider a path *x* in *T* giving rise to the persistence interval [*d*(*R, b*(*x*)), *d*(*R, d*(*x*))]. Then the path *x* will be present in *T*_*r*_ for *r* ≥ *d*(*R, d*(*x*)). However it may happen that the path *x* contains points that are further away from *R* than *d*(*R, b*(*x*)) and will be cut in those points by *B*(*R, r*). This will happen when the path *x* turns around as presented in the Figure 3b. In that instance, the interval [*d*(*R, b*(*x*)), *d*(*R, d*(*x*))] in *T*_*r*_ for certain values of *r* will have a larger value of the first coordinate than the corresponding interval in *T*. Therefore, while the first (birth) coordinate of the interval corresponding to *x* in *T*_*r*_ may be longer than the interval corresponding to *x* in *T*. The actual length can be obtained from the coordinates of degree-2 vertices in *x*.

Those two observations allows for quick computation of the Sholl version of TMD descriptor, i.e. a TMD of a tree *T*_*r*_, also denoted as *TMD*(*T, r*). Firstly the TMD-path decomposition *P* of *T* is computed. Subsequently, for a given radius *r*, a subset *P′* ⊂ *P* containing all the paths *x* such that *d*(*x*) ≤ *r* is selected. The paths in *x* ∈ *P′* are transversed to find the point *f*_*x*_ in there which is inside *B*(*R, r*) and furthest away from its center. Once found, the pair (*d*(*R, f*_*x*_), *d*(*x*)) is added to the *TMD*(*T, r*).

The algorithm described above uses the radial distance from the soma to construct *T*_*r*_. When an intrinsic distance is used instead both in TMD and in construction of *T*_*r*_, the Sholl version of the descriptor is even easier to obtain, as each path *x* ∈ *P* such that *d*(*x*) ≤ *r* will give rise to a pair (*d*(*x*), min(*b*(*x*), *r*)) in *TMD*(*T, r*).

Unlike other Sholl descriptors, *TMD*(*T, r*) has the range in the space of persistence diagrams which is a much richer mathematical structure than real numbers. Yet, it is still possible to compute distances between the functions *TMD*(*T, r*) and *TMD*(*T′, r*). Let us assume that both functions has been computed for a discrete set of values 0 = *r*_0_ *< r*_1_ *< r*_2_ *<* … *< r*_*n*_. Then a distance between *TMD*(*T, r*) and *TMD*(*T′, r*) can be approximated by:

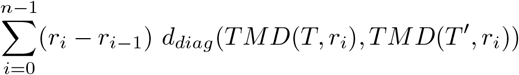

where *d*_*diag*_ denotes any distance between persistence diagrams, e.g. p-Wasserstein distance.

## 2. Stability

In this subsection we define stability and then verify that all Sholl descriptors are stable. We address this issue by verifying that our descriptors are reasonably sensitive to small perturbations of input neurons. More precisely, if two reconstructions of the same neuron vary slightly, they will result in different tree representations. A descriptor is “stable” if, when applied to either tree, it gives results that also vary slightly (i.e. the variation is controlled).

**Fig. 4.**
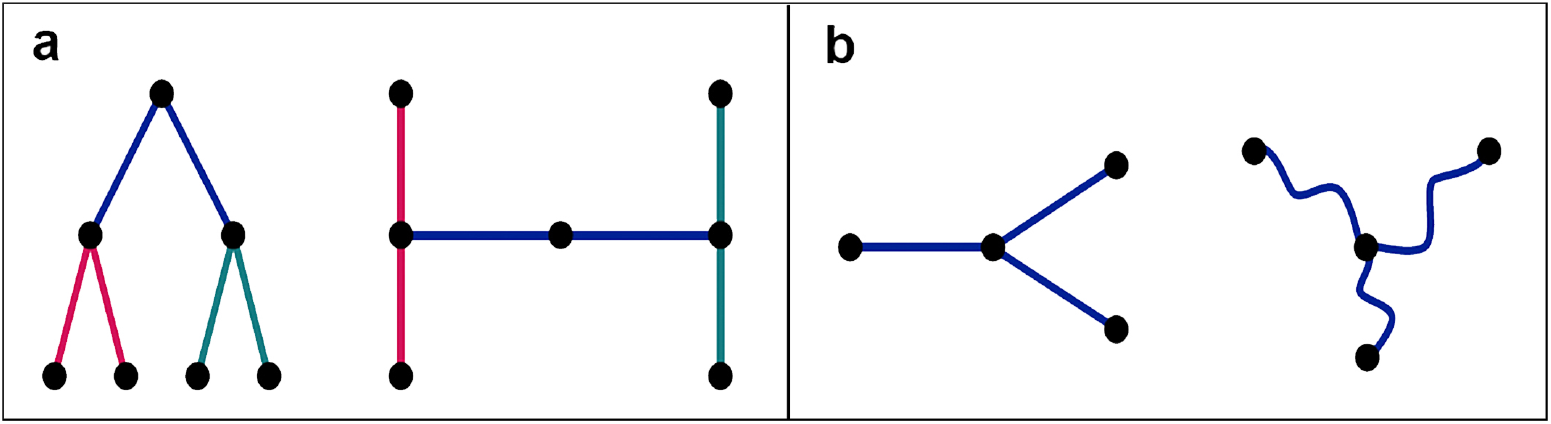
Representative isomorphic trees with entirely different (**a**) branching pattern and tortuosity descriptors.

Two different reconstructions of the same neuron produce two different trees embedded in ℝ^3^. All reasonable reconstruction schemes should produce isomorphic trees, resulting in the same number of primary dendrites and the same number of bifurcations. We can measure the distance between two reconstructed trees under the Hausdorff metric and use it as a measure of closeness. Two such reconstructions are expected to be close in the Hausdorff metric, requiring that our descriptor depends “continuously” on this metric. However, this is not a good notion, as trees that are very close in the Hausdorff metric may still have very different morphological properties (like lengths of branches, number of nodes, etc). Figure 5 gives an illustration of two trees (**b**) and (**c**) close, in a Hausdorff sense, to the initial tree (**a**) note the tree in (**a**) is represented in gray and overlapping with the tree in (**b**), and is depicted below the main branches in (**c**). Clearly the trees (**b**) and (**c**) have distinct morphological features compared to the tree (**a**) and they should not be considered similar.

**Fig. 5.**
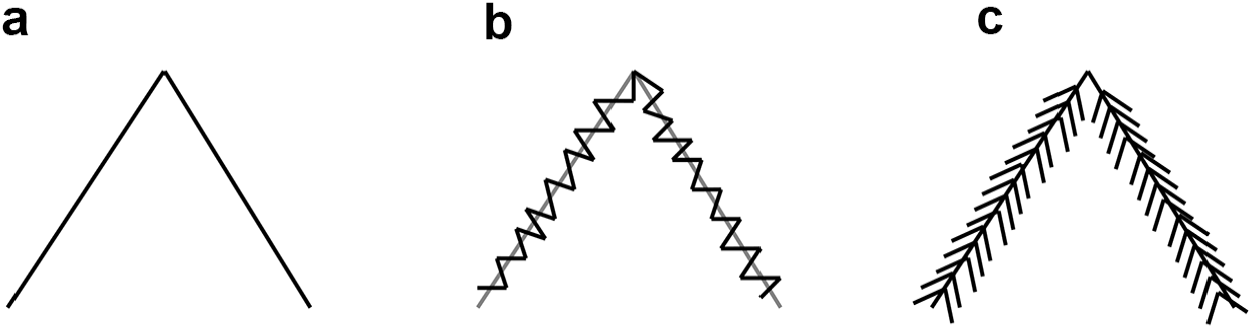
Representative tree (**a**) and similar trees (**b**) and (**c**) that are close to tree (**a**) in the Hausdorff metric.

Our next definition is adapted from ([46], §2) who utilizes it for rectifiable curves and in the context of knot theory. We will assume that the dendrites are piecewise smooth paths in ℝ^3^; meaning that the branches between nodes can be parameterized as *C*^1^-differentiable paths in space.

Again we represent neurons as embedded trees in ℝ^3^. We say that two neurons *N* and *N′* are (*δ, θ*)–close if *N′* can be obtained from *N* by a smooth 1-1 map Ψ supported on an open neighborhood of *N* so that corresponding points *x* and Ψ(*x*) are within *δ* and the norm differences ||*v* − *d*Ψ_*x*_(*v*)|| *< θ* for all *x* ∈ *U* and *v* ∈ *T*_*x*_ℝ^3^, where *d*Ψ_*x*_ is the differential of Ψ at *x*. We recall this is a linear map between tangent spaces *d*Ψ_*x*_ : *T*_*x*_ℝ^3^ → *T*_Ψ(*x*)_ℝ^3^ mapping a vector *v* to *Jac*_*x*_(Ψ)(*v*), where *Jac*_*x*_(Ψ) is the 3 × 3 Jacobian matrix of partial derivatives evaluated at *x*. Let’s now make this construction a bit more precise. For the sake of simplicity we will assume Ψ is defined on all of ℝ^3^.

### Definition 2.1.

We say that *N* and *N′* are (*δ, θ*)-close if there exists an ambiant diffeomorphism Ψ : ℝ^3^ →ℝ^3^ such that |Ψ(*x*) − *x* | *< δ* for all *x*, and the Frobenius norm ||*I* − *Jac*_*x*_Ψ || *< θ* for every *x* ∈ℝ^3^. In contrast with the definition in [46], we not only require the angles between corresponding vectors to be close, but also their norms. This is precisely the essence of the inequality ||*I* − *Jac*_*x*_Ψ|| *< θ*.

### Remark 2.2.

Let’s just observe that our definition is related to the *C*^1^-topology of functions in the following way. If we view a branch *γ* of *N* as a smooth path [0, 1] →ℝ^3^, then it is (*δ, θ*)-close to Ψ ◦ *γ* if both paths are *C*^1^-close. The definition of *C*^1^-closeness doesn’t involve the existence of Ψ, but it cannot be defined globally on neurons, as opposed to just branches, since neuronal trees are not manifolds.

We now define “stability”.

### Definition 2.3.

A Sholl descriptor *ϕ* is *stable* if for any *ϵ >* 0, there exists *η >* 0 so that for *δ < η, θ < η*,

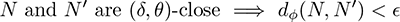

According to this definition, a small perturbation or deformation of the neuron which “moves the points by as little as *δ*” and “distorts the angles by as little as *θ*”, yields a small change in the descriptor *ϕ*.

### Remark 2.4.

Since in this paper we do not distinguish between choices of embeddings of neurons in ℝ^3^, up to *O*(3)-isometry, our definition of stability should operate on isometry classes of neurons. Indeed this can be done as follows. Let [*N*] be the isometry class of *N*, which means that [*N*] is the orbit of *N* under the action of *O*(3), i.e.[*N*] ={*AN, A* ∈ *O*(3)}. We say that [*N*] and [*N′*] are (*δ, θ*)-close if there are representatives *N*_1_ ∈ [*N*] and 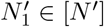 that are (*δ, θ*) close as in Definition 2.3. In our derivations below, when we say that *N* and *N′* are (*ϵ, θ*)-close, we are already thinking of them as being some chosen representatives from their respective isometry classes. It’s easier notation-wise to work with neurons directly rather than with their classes.

Let *N* be a neuron represented as a spatial tree, and let *ϕ* be a Sholl descriptor. The nodes for *N* are sorted according to increasing distances from the soma 0 *< r*_1_ *<…< r*_*k*_, with *k* being the number of nodes. These distances are radial or dendritic depending on the descriptor. We make the assumption that a deformation of a neuron does not introduce new bifurcations, and so the leaf index is completely unchanged by deformation. It is evidently stable.

We start by verifying the stability of the branching pattern descriptor. This descriptor is only based on the distribution of nodes, and so only the *δ* constant matters. The nodes have (normalized) distances from the soma *r*_*i*_ ≤ 1. Let’s move the nodes of *N* by a distance *δ*, and by that we mean there is a homeomorphism Ψ : *N* → *N′*, taking node *P*_*i*_ to 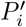 so that for all *i*, 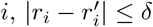. Note that we start with *N* that is normalized (i.e. it is within a ball of radius 1) but then *N′* might not be. By normalizing *N′* to *N ″*, and taking the new node radii 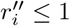, we see that

**Fig. 6.**
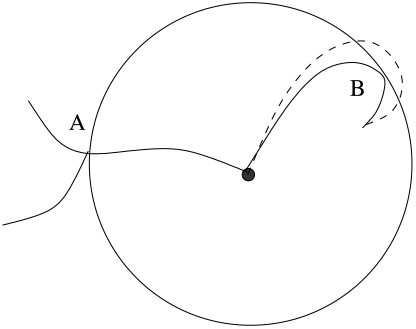
Instability behavior for tortuosity descriptor.

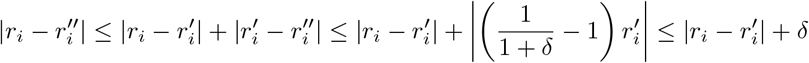

So we can assume from the start that both nodes of *N* and *N′* are such that *r*_*i*_, 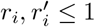, and that for all *i*, 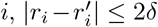. Recall next that the *t*_*i*_’s are a renaming of the *r*_*i*_ and 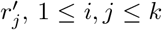, 1 ≤ *i, j* ≤ *k*, once they are ordered by increasing order, so that *t*_1_ *< t*_2_ *<* … *< t*_2*k*_ (see (4)). Given this, the formula for functional distance takes the form

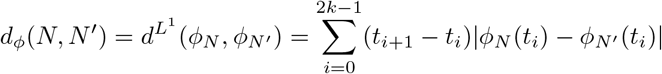

Choose *δ <* min_*i*_{|*r*_*i*_ − *r*_*i*+1_|}. By construction of the branching pattern, the difference *ϕ*_*N*_ (*t*_*i*_) − *ϕ*_*N′*_ (*t*_*i*_) can be at most ±1 on intervals [*t*_*i*_, *t*_*i*+1_[of the form 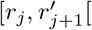 or 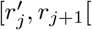 for some *j* (*j* need not be same as *i* since this is obtained after renumbering), and is zero otherwise. These intervals are of length 2*δ* or less. This gives that *d*_*ϕ*_(*N, N′*) ≤2*k* 2*δ* = 4*kδ*, where *k* is the number of nodes. By choosing *δ* small enough, this is less than any desired *ϵ*.

The taper and energy descriptors depend only on the node distribution, and are thus stable thanks to the arguments presented so far. Note that for the taper rate, we assume that the width of a dendrite at a given node is the same in any reconstruction (this is not a varying feature in our definition of stability), so this descriptor only depends on nodes, and it is stable.

To see stability of the TMD with the radial distance to the soma let us observe that the TMD-path decomposition may change when the position of nodes is perturbed. This will happen when the endpoints of two paths that merge in a bifurcation point *b* are at almost the same distance from the soma. In this case a perturbation of the endpoints of those paths may result in swapping the branch that continues up from *b* with the one that terminates there. However, since the TMD only gathers the values of distances from the soma, the endpoints of persistence intervals will move by at most *δ* which directly translate in stability of the descriptor.

As for the Tortuosity descriptor, we recall that *T* associates to every *N* and *r* ∈ [0, 1] the average tortuosity of *N*_*r*_ which is the connected component of *N* containing the soma inside the ball of radius *r*. The stability for *T* holds under one condition, and so we refer to it as “conditional stability”. We discuss this condition next and remark that it is almost always realized so that in practice and generically *T* behaves stably. To understand this condition, we must observe that instability can occur in the following situation illustrated in Figure 6.

Here *r* is chosen to be the radial distance of the node *A*. The twisted dendrite touches the sphere tangentially at *B*. When measuring tortuosity of *N*_*r*_, we only consider the term *δ*_*SB*_ (6) which is the tortuosity from the soma *S* to *B* considered as a leaf. If a small perturbation causes the branch to be inside the sphere, this term is replaced by the term *δ*_*SC*_ which is the tortuosity of the entire branch inside *B*(*r*) from *S* to *C*. This leads to a sudden increase in tortuosity which can potentially lead to instability. However this only happens if indeed a sphere through a node is tangent to a branch, which is a rare instance.

Let *γ* : [0, 1] **→** ℝ^3^ be a smooth space curve. Then its length is given by 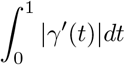. We say that the two paths *γ*_1_ and *γ*_2_ are (*δ, θ*) close if there is a (*δ, θ*)-diffeomorphism taking *γ*_1_ to *γ*_2_; that is *γ*_2_ := Ψ ◦ *γ*_1_.

### Lemma 2.5.

*Let γ*_1_ *be a given smooth curve in* ℝ^3^. *For every ϵ >* 0, ∃*η with δ < η, θ < η, such that for every curve γ*_2_ *that is* (*δ, θ*)*-close to γ*_1_,

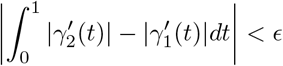

*Proof*. Let *γ*_2_ be (*δ, θ*)-close to *γ*_1_, meaning there is a (*δ, θ*)-diffeomorphisme Ψ taking *γ*_1_ to *γ*_2_. Since *γ*_1_ is differentiable, 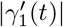 is bounded uniformly on [0, 1], say by *M >* 0. We can write

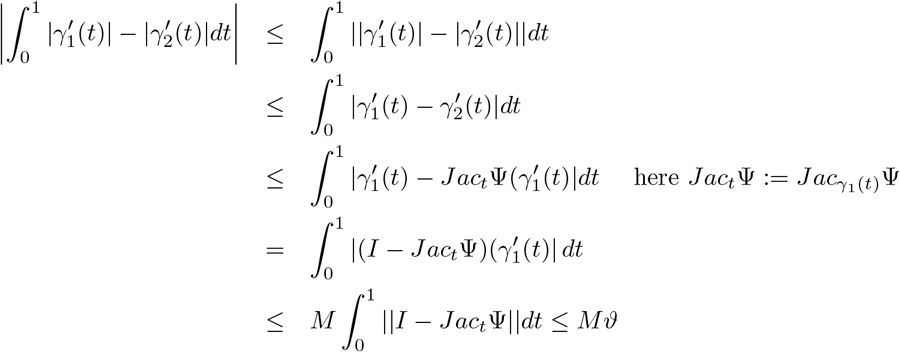

where we know by definition that ||*I* − *Jac*_*t*_Ψ|| ≤ *θ* on the domain of Ψ. By choosing 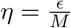, we obtain our claim. □

Let Ψ be (*δ, θ*)-diffeomorphism taking *N* to *N′*. The map Ψ maps nodes to nodes, branches to branches necessarily. Lemma 2.5 shows that by controlling (*δ, θ*) we can control the lengths of branches. It is clear that *T* is stable away from the stated condition, meaning that if spheres through the nodes of *N* and *N′* are not tangent to dendrites, then *d*_*T*_ (*N, N′*) = *d*_*T*_ (*N*, Ψ(*N*)) *< ϵ* for any chosen *ϵ >* 0, once *δ, θ* are chosen sufficiently small.

Finally we discuss the stability of the flux descriptor. The stability in these cases hinges on controlling the variation of angles in any neuron deformation. This is a direct consequence of the following lemma.

### Lemma 2.6.

*Let v*_1_, *v*_2_ ∈ *T*_*x*_ℝ^3^ *be unit vectors, x* ∈ *N (x will be typically a node in our case). Then for any ϵ >* 0, ∃*η >* 0 *such that for any* (*δ, θ*)*-diffeomorphism* Ψ *with θ < η*,

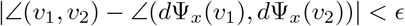

*Proof*. We fix *x* and drop it from the notation. We write 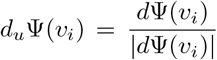 the normalized vector. Using the cosine-angle formula |*v*_1_ − *v*_2_|^2^ = |*v*_1_|^2^ + |*v*_2_|^2^ − 2|*v*_1_||*v*_2_| cos ∠(*v*_1_, *v*_2_), we can write

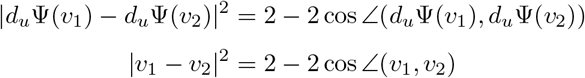

By taking the difference, we see immediately that we can make the angles ∠(*v*_1_, *v*_2_) and ∠(*d*Ψ_*x*_(*v*_1_), *d*Ψ_*x*_(*v*_2_)) arbitrarily close by making their cosines arbitrarily close or equivalently by making |*d*_*u*_Ψ(*v*_1_) − *d*_*u*_Ψ(*v*_2_)|and |*v*_1_ − *v*_2_|arbitrarily close. We check this last part.

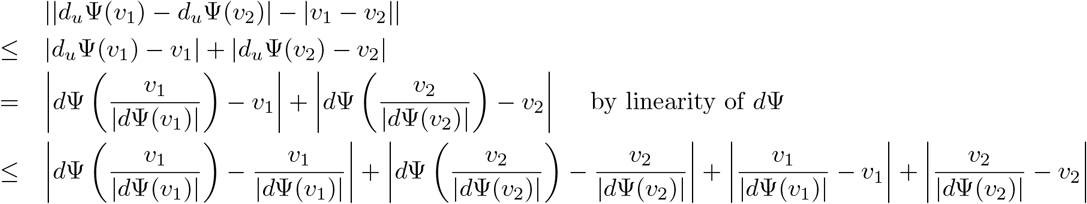

Since for any vector *v* ∈ *T*_*x*_ℝ^3^, |*d*Ψ(*v*) − *v*| *< θ*, we can extend the string of inequalities above to

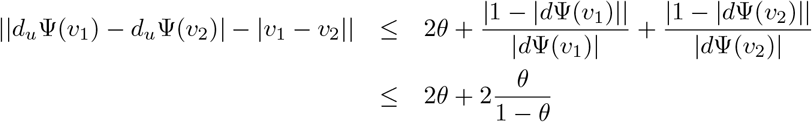

By making *θ* small, the left term can be made arbitrarily small and this is enough to yield our claim. □

## 3. The Detection Algorithm

The detection construction and motivation was discussed in §1.4. In this section we define and compute the level of detection of a class of neurons, within a collection, by a Sholl descriptor *ϕ*.

Let 𝒩 = {*N*_1_, …, *N*_*k*_} be a set of neurons divided up into classes. A neuron *N*_*i*_ will belong to a class *C*(*N*_*i*_).Recall that associated to *ϕ* we have a pseudo-metric *d*_*ϕ*_ on the set of neurons (2).

For every neuron *N* ∈ {*N*_1_, …, *N*_*k*_}, let *σ* be the permutation of {1, …, *k}* that orders the neurons according to their distance from *N*, so that

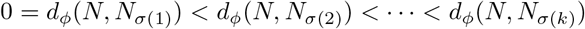

- For every 1 ≤ *i* ≤ *k*, let *det*_*ϕ*_(*N, i*) denote the smaller of the two numbers:

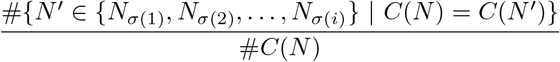

and

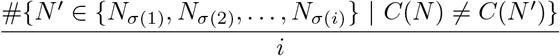

where # means cardinality of set.
- The local detection rate at *N, det*_*ϕ*_(*N*), is defined as the maximum value of *det*_*ϕ*_(*N, i*), for 1 ≤*i* ≤*k*.
- The detection rate of a class *C* is now equal to the maximum of all local detection rates of neurons that belong to that class. Formally

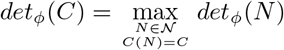

We suspect the existence of other machine learning procedures that capture the same percentiles. We are not aware of any reference however.

### Example 3.1.

Suppose det_*ϕ*_(*C*) = 75%. This implies there is a ball in the *d*_*ϕ*_ metric that contains (at least) 75% of all the internal neurons in *C*, and within that ball, (at least) 75% of all neurons are from *C*. A detection rate of 100% means perfect detection whereby there is a ball containing all of *C* and no other neurons from any of the other classes.

## 4. Combination of Descriptors and Classification

Our objective is to build a collection of distances *d*_1_, …, *d*_*n*_ that can be used to discriminate among trees (classes of neurons) according to a given morphological feature. One aim is to understand, for two or more classes of trees, which morphological features differentiate them. For example, *C*_1_ can be a class that represents neurons from an experimental group with a neurological disease, while *C*_2_ is a class of neurons from a control group.

### 4.1. Vectorization

The following vectorization method is used for a purpose of unsupervised classification. The starting point is a set *C* = {*N*_1_, …, *N*_*k*_}of neurons and a collection of descriptors. The values of the considered descriptor at 0 and 1 are meaningful and will be entries of the constructed vector. In particular those values for a given neuron *N* are:

- *B*_*N*_ (1) for branching, being the (opposite) of the number of primary branches.
- *To*_*N*_ (1) for tortuosity, being the average tortuosity of *N*.
- *Li*_*N*_ (0) for leaf index, being the total number of leaves of *N*.
- *E*_*N*_ (1) for energy, representing the total energy vector at the soma.
- *F*_*N*_ (1) for flux, representing the total flux vector at the soma.
- *Tr*(1) for taper rate, representing the total taper rate vector at the soma.
- *W*_*N*_ (1) for wiring, representing the total wiring vector at the soma.

Given a descriptor *ϕ*, being one of the descriptors above, and a neuron *N*, we also consider its area value 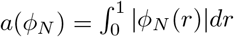.

We now list all of our descriptors associated to the neuron *N* in the order as above. The corresponding vector in ℝ^14^ is:

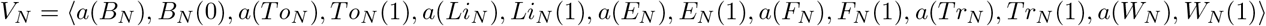

If a descriptor data is not available (eg. taper rate), then the corresponding pair of entries is omitted from the vector. For Dataset 1, we vectorized based on seven descriptors (all but TMD-Sholl and taper rate).

We want to highlight that this is just one of possibly many ways to vectorize real-valued functions. Another vectorization method, suitable for labeled data, is presented in the Section S4.3. Lastly, when our Sholl function happens to have values in more general metric space (as in case of Sholl-TMD that has values in a space of persistence diagrams), we propose to select a grid of values of the radi and, for each of them, vectorize the given persistence diagram.

### 4.2. Combination of Metrics

Let us assume that the considered collection of neurons belongs to a number of classes. We present a greedy grid–based search procedure that can reveal which features are ‘fundamentally’ different between the classes. For that purpose, we consider a new distance *d* being the following linear combination:

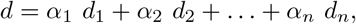

where the non-negative constants *α*_*i*_ are sampled from a uniform grid.

Consider first the situation when each neuron may belong to one of two classes. For the fixed choice of *α*_1_, …, *α*_*n*_ let 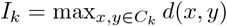 be the maximal distance *d* between objects in *C*_*k*_ for *k* ∈{1, 2}. Let 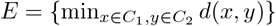 be the minimal distance *d* between elements from two different classes. We then consider the ratio

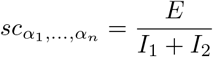

and select *α*_1_, …, *α*_*n*_ from our grid of points that maximize 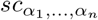. Note that this greedy grid search has exponential complexity with *n*, being the number of considered distances.

The obtained weights *α*_1_, …, *α*_*n*_ give an idea of the relative importance of different distances in the separation of classes *C*_1_ and *C*_2_. Consequently, when features leading to distances *d*_1_, …, *d*_*n*_ can be interpreted geometrically, the weights may help identify the geometrical features that are important in the separation of the classes, and those that are not.

This idea can be generalized for a multi–class problem. There are various ways this can be achieved and we will present one possible approach. Let us have *k* classes *C*_1_, …, *C*_*k*_ and define 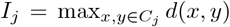 (maximal internal distance in *C*_*j*_) and 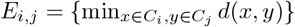 (minimal distance between classess *C*_*i*_ and *C*_*j*_). Then the multi-class score is given by:

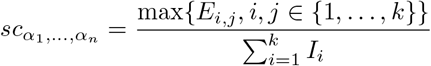

The remaining of the procedure described above is not changed. We can show that separation is meaningful, and not a result of over-fitting, by conducting a “permutation test” (see §4.4).

**Fig. 7.**
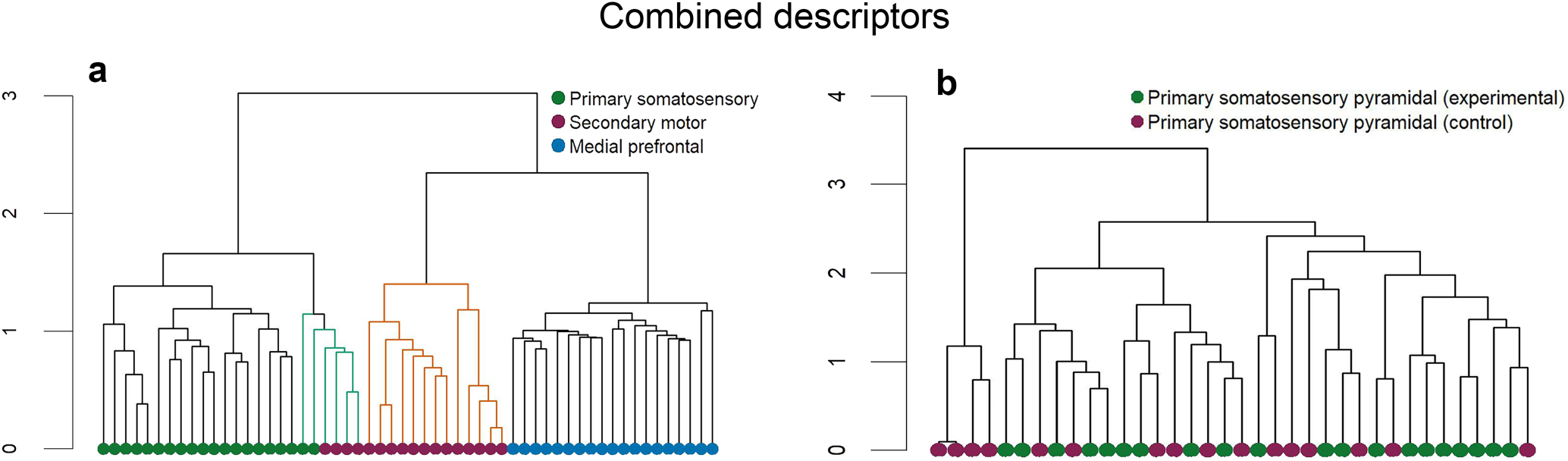
Representative dendrograms based on the combination of metrics.(**a**) Combined dendrogram for Dataset 4 and (**b**) Dataset 5.

### 4.3. Metric Learning

We use Metric Learning for a labeled collection of neurons 𝒩 partitioned into *k*-classes labeled by *ℓ*_1_, …, *ℓ*_*k*_. Every neuron *N* ∈𝒩 has therefore some label *ℓ*(*N*) = *ℓ*_*i*_ for some *i*. Firstly, the input collection of neurons will be vectorized using a number of Sholl descriptors with a label-aware method presented below. Subsequently, the Euclidean metric that can be used to compare the vectorized vectors will be adjusted to separate the classes.

Let Ω = {*ϕ*_1_, …, *ϕ*_*m*_} be the considered family of Sholl descriptors. For *N* ∈ 𝒩, we can consider the average distance for the descriptor *ϕ*_*j*_ of *N* to all neurons labeled by *ℓ*_*i*_;

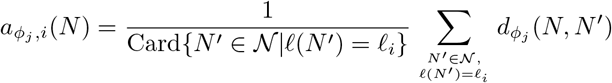

As *ϕ* runs over Ω, we obtain the vector in ℝ^*mk*^ that characterizes the neuron *N* ;

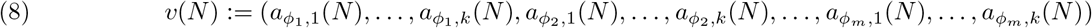

where the first *k* entries give the average distances of *N* to all *k* classes in the metric 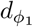, and the last *k* entries give the average distances of *N* to all *k* classes in the metric 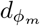. The vector in (8) depends on the ordering on the labels and Sholl descriptors, but the final outcome will not. In all cases, and for the remaining constructions in this section, an order on classes and on descriptors *ϕ*_1_, …, *ϕ*_*m*_ is always chosen before we start running any algorithm, and this order is preserved throughout the process.

Starting with a dataset of neurons of a cardinality *n*, and given *m* descriptors, we obtain a set of *n* labeled vectors in ℝ^*mk*^, one for each neuron. The obtained vectors will be labeled in accordance to the labels of neurons.

Ideally one would hope that the Euclidean metric 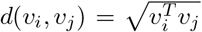 does “separate” the classes, meaning that vectors in the same class are close and those in different classes remain relatively distant. This is hardly the case in practice, so one seeks a modification of this Euclidean metric which has this separation property. A standard approach is to introduce a “matrix of weights” *M*, which is *k* × *k*, positive, so that 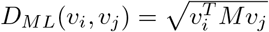 defines a new metric on ℝ^*n*^ (so called Mahalanobis metric, see [45]) with better separating properties with respect to the chosen classes. More precisely, *D*_*ML*_ maximizes the sum of distances between points with different labels while keeping the sum of distances between those with similar labels small. Note that since *M* can be written as *LL*^*T*^, the associated *D*_*ML*_ metric has the following interpretation: it is the distance obtained by first moving vectors via *L* in ℝ^*m*^*k*, then taking their Euclidean distance. Various machine learning algorithms have been implemented to find this optimal *M*. This approach is entirely supervised since we need the classes to train the matrix entries and thus the metric.

This vector of points in ℝ^*mk*^ is a desired input for the Metric Learning procedures implemented for instance in [31]. The vectorized data are therefore run through a supervised ML algorithm [31]. The end result is a metric *D*_*ML*_, depending on 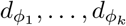, which differentiates between the classes of neurons.

There are many ways in which the obtained metric can be used. For instance, given a new neuron *N*, we may wish to know how close in feature is *N* to the classes of neurons considered in 𝒩. For that purpose, we may apply the following procedure;

- Run metric learning on the vectorized classes of 𝒩 to obtain a new metric *D*_*ML*_. The new metric is validated after being tested for “overfitting” (see §4.4). A good metric gives good separation of the vectorized classes.
- Given a neuron *N*, vectorize it as in (8) and then take its average distances to the classes in 𝒩.
- Pick the class that is, on average, closest to the new neuron *N*.
- (feature selection) Run each descriptor on the classes. If the detection rates are lower than 80% on all classes, the descriptor can be considered “noisy” and is subsequently excluded. Repeat the process above with non noisy descriptors.

### 4.4. Overfitting

Both grid search §4.2 and metric learning §4.3 provide efficient tools to differentiate classes of neurons. Yet, the fact that these methods return clear separation between two or more classes of trees is not sufficient to conclude that the separation is geometrically meaningful. By itself, those methods may fir exactly against the data.

#### Example 4.1.

Let us consider four vertices of a square: *A* = (1, 1), *B* = (−1, 1), *C* = (−1, − 1) and *D* = (1, − 1). Suppose that the first class consists of points *A* and *D*, while the second class is composed of points *B* and *C*. Metric learning, for instance large margin nearest neighbor algorithm we use, will seek to place elements of different classes far away and those of the same class close together. This can be achieved by a metric function 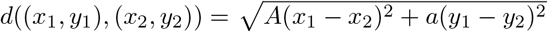 by making *A* large and *a* small. Such a perturbed Euclidean metric can be obtained both by the grid search and metric learning. Yet, it is clear that the division to the first and second class is somewhat arbitrary; In fact, putting points *A* and *B* to the first class and *C* and *D* to the second one is geometrically similar. Alternatively, separation of these classes can be achieved by the same distance function with *A* small and *a* large, and therefore can also be found by the methods we present here.

There are two ways an overfitting, as the one described in Example 4.1 may be detected;

First, when the dataset is large enough, a standard k-fold cross-validation is applied. In this case, a method will be repetitively trained on a subset of the considered dataset, and tested on the remaining test set. Once the results obtained on the test sets are good, we can assume that the methods do not overfit.

In the second case, which is used for small datasets, a procedure similar to a *permutation test* is used. Namely, after obtaining separation of the given classes, we will repetitively permute all the labels of data points and run the grid search / metric learning for the data with the permuted labels. We will check how frequently a good separation between the permuted labels is obtained. If that happens often, then the separation between the initial classes is not valid, as it is rather a result of overfitting. However, of it is not the case, then we have additional evidence that the separation of the original classes is meaningful.

## Footnotes

1 Consequently, all branchpoints in a neuronal tree, with exception of the soma, have degree two.

2 In case of two or more paths satisfying this condition, the one that continue is picked up randomly.

